# Polygenic local adaptation in metapopulations: a stochastic eco-evolutionary model

**DOI:** 10.1101/2020.06.16.154245

**Authors:** Enikő Szép, Himani Sachdeva, Nick Barton

## Abstract

This paper analyses the conditions for local adaptation in a metapopulation with infinitely many islands under a model of hard selection, where population size depends on local fitness. Each island belongs to one of two distinct ecological niches or habitats. Fitness is influenced by an additive trait which is under habitat-dependent directional selection. Our analysis is based on the diffusion approximation and accounts for both genetic drift and demographic stochasticity. By neglecting linkage disequilibria, it yields the joint distribution of allele frequencies and population size on each island. We find that under hard selection, the conditions for local adaptation in a rare habitat are more restrictive for more polygenic traits: even moderate migration load per locus at very many loci is sufficient for population sizes to decline. This further reduces the efficacy of selection at individual loci due to increased drift and because smaller populations are more prone to swamping due to migration, causing a positive feedback between increasing maladaptation and declining population sizes. Our analysis also highlights the importance of demographic stochasticity, which exacerbates the decline in numbers of maladapted populations, leading to population collapse in the rare habitat at significantly lower migration than predicted by deterministic arguments.

## Introduction

Adaptation to environmental change is often rapid (Thompson, 1998; Grant and Grant, 2006; Kokko and López-Sepulcre, 2007; Kinnison and Hendry, 2001) and time scales of ecological and evolutionary change comparable, giving a feedback between demography and evolution. Fisher (1930) first described reciprocal interactions between population size and adaptation, leading to the notion of hard selection, whereby high genetic load drives populations to extinction. This is an extreme example of a more general feedback: an increase in genetic load due to deleterious variants reduces population size; smaller populations are affected more strongly by drift and gene flow, which increase the fixation of locally deleterious alleles, further decreasing size.

Such eco-evolutionary feedbacks are crucial during evolutionary rescue following a sudden environmental shift (Gomulkiewicz and Holt, 1995; Gonzalez et al., 2013), and are key to the survival of marginal populations (Kawecki, 2008), the colonisation of peripheral habitats (Barton and Etheridge, 2018; Sachdeva, 2019), and the emergence of sharp range margins in the absence of environmental discontinuities (Polechová and Barton, 2015; Polechová, 2018).

Eco-evolutionary feedbacks are especially important in fragmented habitats, where stochastic extinction and recolonisation of patches requires recurrent bouts of rapid adaptation, especially if selective pressures vary across patches. This kind of metapopulation structure may arise, for instance, if multiple hosts are available within the same region (Carroll and Boyd, 1992; Dobler and Farrell, 1999), favouring host-specific adaptations. The potential for local adaptation and the stability of sub-populations then depends on the interaction between selection (which is mediated by the genetic architecture of selected traits), dispersal (which protects populations from inbreeding load and stochastic extinction, but may also introduce maladapted phenotypes) and demography (which is affected by mean genetic fitness, and in turn influences the efficacy of selection).

Previous theory on the persistence of subdivided populations neglects key aspects of this interplay. For instance, Blanquart et al. (2012) analyse conditions for local adaptation in a spatially heterogeneous metapopulation under *soft* selection (i.e., constant population sizes), thus neglecting the feedback between fitness and demography. Ronce and Kirkpatrick (2001) explicitly model the effects of maladaptation on population sizes in a metapopulation with multiple ecologically distinct habitats, but neglect all stochasticity. Another approach, exemplified by Hanski and Mononen (2011), considers re-colonisation of patches to be instantaneous and extinction to depend deterministically on patch fitness. However, this fails to address how the coupled *stochastic* dynamics of genotype frequencies and population sizes influence extinction thresholds. Conversely, almost all work on stochastic fluctuations in metapopulations neglects natural selection (Lande, 1993; Lande et al., 1998; Ovaskainen and Meerson, 2010).

A better understanding of the link between maladaptation and extinction also requires genetically realistic eco-evolutionary models. However, at present, most metapopulation models assume one of two extreme architectures (Govaert et al., 2019; Lion, 2018): either mutations occur one by one (adaptive dynamics), or there are very many infinitesimal-effect loci (quantitative genetics). While the latter can describe response from standing variation (which often underlies rapid adaptation), with a few exceptions (Hanski and Mononen, 2011; Ronce and Kirkpatrick, 2001), quantitative genetic models only deal with migration into a *single* population (e.g., Barton and Etheridge (2018); Chevin et al. (2017); Tufto (2001)).

Here, we investigate the joint evolution of population size and allele frequencies in a metapopulation consisting of infinitely many islands connected via migration. Each island belongs to one of two different habitats characterised by distinct selection pressures. We ask: when are demographically stable, locally adapted populations maintained within islands belonging to the ‘rare’ or marginal habitat? Conversely, when does maladaptive gene flow from the abundant habitat reduce the rare habitat to a maladapted (and possibly nearly extinct) sink? Understanding evolution in marginal habitats has important implications for range limits and the long-term survival of metapopulations; moreover, local adaptation in marginal habitats may be the first step towards speciation.

A key focus of our work is on how the coupling between population size and mean fitness (‘hard selection’) influences local adaptation and extinctions. Such coupling places severe constraints on the survival and adaptation of interconnected populations: gene flow limits local adaptation not only by overwhelming selection at individual loci, but also through the effect of migration load on population size. Segregation of locally maladaptive alleles at many loci at even low frequencies substantially reduce mean fitness, causing lower population numbers, which further impairs selection at individual loci, resulting in a positive feedback between population decline and loss of local adaptation (‘migrational meltdown’; see Ronce and Kirkpatrick (2001)).

As we demonstrate below, random fluctuations in population size (demographic stochasticity) as well as in allele frequencies (genetic drift) strongly influence thresholds for maladaptation and extinction. Thus, going beyond deterministic analyses by considering both sources of stochasticity within a common framework is crucial for understanding extinction: smaller populations are more prone to fix maladaptive alleles due to genetic drift and swamping from larger populations; this further reduces the fitness and size of small populations, rendering them even more vulnerable to demographic fluctuations and chance extinction, even in parameter regimes where demographic stochasticity *by itself* (i.e., in the absence of maladaptation) is of little consequence.

A second focus is to understand how the feedback between population size and fitness is mediated by the genetic basis of selected traits. In particular, is maladaptation (and possibly extinction) in the rare habitat more or less likely for more polygenic traits? Understanding polygenic local adaptation within subdivided populations is challenging due to statistical associations, i.e., linkage disequilibria (LD) between loci. Our theoretical analysis neglects such associations, assuming linkage equilibrium (LE) within demes, and becomes exact in the limit where all processes are much slower than recombination. Assuming LE allows us to work solely with allele frequencies. However, LE does *not* imply that loci evolve independently: under hard selection, evolutionary dynamics of different loci become coupled through their aggregate effects on population size, which in turn influences individual loci via genetic drift.

Our analysis is based on a *diffusion approximation* for the joint stochastic evolution of allele frequencies and population size. The diffusion approximation has been widely used in population genetics (Fisher, 1922; Kimura, 1955), but remains less prominent in ecology, and has only been used to model stochastic population dynamics, without genetics (e.g., Lande (1993); Mangel and Tier (1993)). Our framework incorporates both demographic stochasticity and genetic drift, following Banglawala (2010). The full model requires numerical solution, but explicit analytical predictions are possible in various biologically interesting limits. In order to assess the importance of LE and other assumptions, we compare theoretical predictions against individual-based simulations with a finite number of demes.

## Model and Methods

Consider a metapopulation with infinitely many islands (demes) that exchange genes via a common pool. We assume that any island belongs to one of two local habitats, indexed by *α*=1, 2. A fraction 1 − *ρ* (or *ρ*) of islands belong to the first (or second) habitat. We choose *ρ*<1*/*2, such that *ρ* always denotes the fraction in the rare habitat, indexed as habitat 2 hereafter.

Individuals are haploid and express an additive trait influenced by *L* unlinked, biallelic loci with alternative states denoted by *X*=0, 1. The trait is assumed to be under directional selection towards one or the other extreme: thus, a genotype with all ‘1’ alleles or all ‘0’ alleles has maximum fitness in the first or second habitat respectively. For simplicity, the maximum possible genetic fitness in either habitat is assumed to be the same. Thus, individual fitness is given by exp 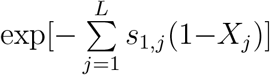 and exp 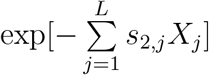 in the two habitats, where *s*_1,*j*_>0 (or *s*_2,*j*_>0) denotes the strength of selection against the locally disfavoured allele at locus *j* in habitat 1 (or 2), and *X*_*j*_ the allelic state at locus *j*. The life cycle of individuals consists of dispersal, followed by selection and mating.

As our focus is on how gene flow influences selected polymorphisms, we neglect other sources of variation. However, the framework easily extends to include mutation. In each generation, individuals migrate with probability *m* into a common pool; migrants from this pool are then evenly redistributed over islands. The assumption of infinitely many islands means that genotype frequencies in the migrant pool are essentially deterministic; in simulations, we model a large but finite number of islands.

We assume hard selection, where population sizes are stochastic but influenced by mean fitness on the island plus local density-dependent regulation: the size 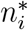 on island *i*, after selection and regulation, is a Poisson random variable with mean 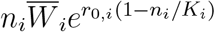. Here, *r*_0,*i*_≥0 is the baseline rate of growth, *K*_*i*_ the baseline carrying capacity, *n*_*i*_ the population size prior to selection, and 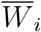 the mean genetic fitness on island *i*. For simplicity, we assume *r*_0,*i*_=*r*_0_ and *K*_*i*_=*K* across all islands. The 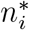 offspring are formed by randomly sampling 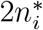 parents (with replacement) from the *n*_*i*_ individuals in proportion to individual fitness, and then creating offspring via free recombination of each pair of parent genotypes. Note that selection is density-independent, i.e., relative fitness of genotypes is independent of population size.

When selection is strong relative to migration and drift, and the number of loci not very large, populations can adapt to their local habitat, resulting in LD between alleles favoured within a habitat. Our theoretical framework, however, assumes that selection per locus and migration are weak compared to recombination, such that LD within a deme (generated by immigration of individuals from differently adapted habitats) is rapidly dissipated and can be neglected. This allows us to consider the coupled dynamics of population size and *L* alleles (rather than 2^*L*^ genotypes) while still accounting for stochastic effects via the diffusion approximation.

For weak growth, selection and migration (i.e., *r*_0_, *s, m*≪1), we can use a continuous time approximation for allele frequency and population size dynamics. The size *n*_*i*_ and the allele frequency *p*_*i,j*_ at the *j*^*th*^ locus on the *i*^*th*^ island satisfy the following *coupled* equations (see also Appendix A1, SI for details):

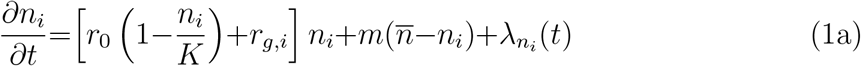

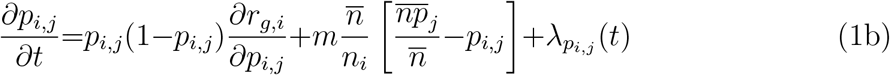

Here, *r*_*g,i*_ is the genetic component of the growth rate (i.e., the log fitness) averaged over all genotypes on island *i*. For the fitness functions described above, we have: 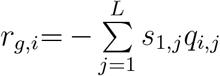 for an island in the first habitat and 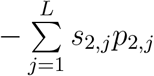 in the second habitat. Here *p*_*i,j*_ and *q*_*i,j*_=1 − *p*_*i,j*_ denote the frequencies of the ‘1’ and ‘0’ alleles at locus *j* on island *i*.

Note that the dynamics of any one deme are coupled to the dynamics of all other demes via the mean number of immigrants 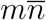 and the mean number of immigrant alleles 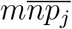 (at locus *j*) per unit time (where 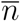 is the population size and 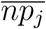 the number of allele copies per deme, averaged over the metapopulation).

Equation (1a) describes how population size evolves over time on an island in a given habitat. The first term within the square brackets describes logistic growth, and the second how growth rates are reduced relative to the baseline *r*_0_ due to habitat-dependent selection against locally disfavoured alleles; this term *r*_*g,i*_ couples population sizes to allele frequencies. The second term describes migration, which makes a net positive contribution when the focal deme is smaller than the average 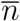 across the metapopulation. The third term 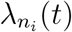 is an uncorrelated random process with 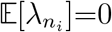 and 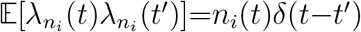, where 𝔼[…] denotes an average over independent realisations. This describes fluctuations of population size due to demographic stochasticity, inherent in reproduction and death. Since the number of offspring is Poisson-distributed, the variance of population sizes is just *n*_*i*_(*t*). Note that the noise term can generate local extinctions even in well-adapted populations: extinction arises from the stochastic dynamics, rather than being imposed arbitrarily.

Equation (1b) describes allele frequency dynamics at locus *j*: the first term corresponds to selection against the locally deleterious allele; the second term describes the effect of migration, which pulls allele frequencies towards the metapopulation average 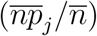, which is the allele frequency in the migrant pool. Islands with larger populations contribute more to the migrant pool; they are also less prone to swamping by maladaptive alleles, since the migration term in eq. (1b) is inversely proportional to *n*_*i*_. This results in a positive feedback: better adapted islands are more populous, send out more migrants and are less affected by incoming, maladapted individuals, maintaining local adaptation more easily. Fluctuations in allele frequencies are described by 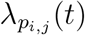, which satisfies 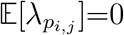 and 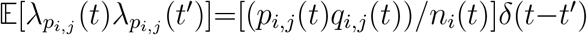 (where *q*(*t*)=1− *p*(*t*)), as in the haploid Wright-Fisher model.

Equations (1a) and (1b) can be made dimensionless by rescaling population size to the carrying capacity *K*, and all evolutionary rates to the baseline growth rate *r*_0_. This gives the following re-scaled parameters (denoted by uppercase letters): *T* =*r*_0_*t, M* =*m/r*_0_, *S* = *s/r*_0_, *N* =*n/K*, and the new parameter *ζ*=*r*_0_*K*, which represents the number of births per unit time at carrying capacity, and hence governs the magnitude of demographic fluctuations. Both scaled and unscaled parameters are tabulated in Key Notation.

### Diffusion approximation for the joint distribution of allele frequencies and population size

The time evolution of the joint probability distribution Ψ_*i*_(*N, p*_1_, …, *p*_*L*_) of allele frequencies {*p*_1_, …, *p*_*L*_} and (re-scaled) population size *N* on any island in habitat *i* is described by the diffusion approximation, which depends only on the mean and variance of the change in *N* and {*p*} per unit time. For ease of notation, the vector (*N, p*_1_, … *p*_*L*_) is denoted by **x**. We have:

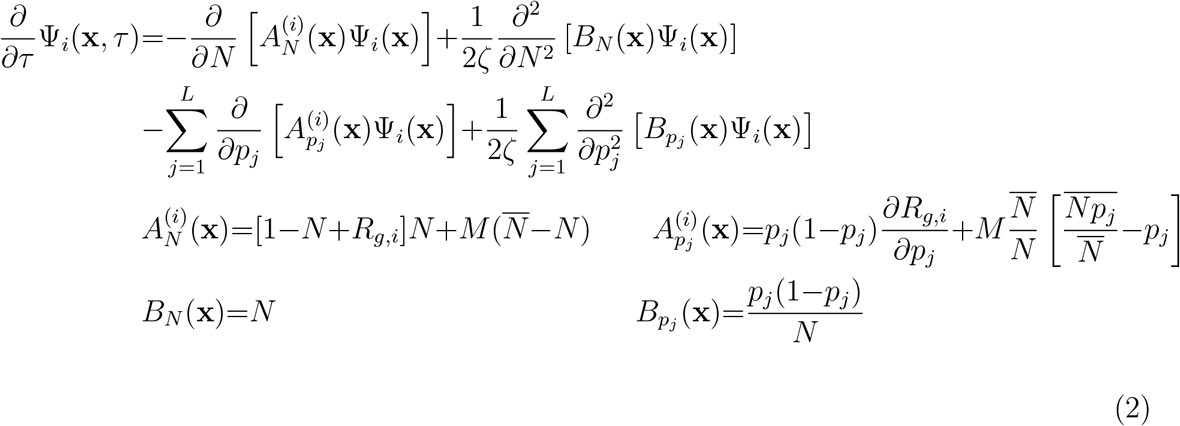

The equation is expressed in terms of re-scaled parameters and *ζ*=*r*_0_*K* (see above). Here, 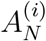 and 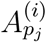 specify the expected rate of change of the population size and allele frequencies on an island in habitat *i* (see also eq. (1)), and *B*_*N*_ and 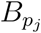 variance of the change per unit time (which is independent of habitat). The dependence on the local habitat arises only through the average log fitness *R*_*g,i*_=*r*_*g,i*_*/r*_0_, given by 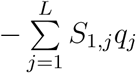 and 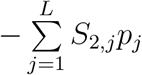 in the first and second habitats respectively.

In principle, equation (2) can be numerically integrated to obtain the joint distribution of *N* and {*p*_*j*_} through time. However, we focus on the stationary (equilibrium) distribution. This depends on the numbers and genetic composition of the migrant pool, which is determined by the average number of individuals 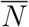 and allele copies 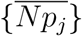 (for each locus) across the metapopulation. The stationary distribution on any island in habitat *i* is given by (see Appendix A2, SI for details):

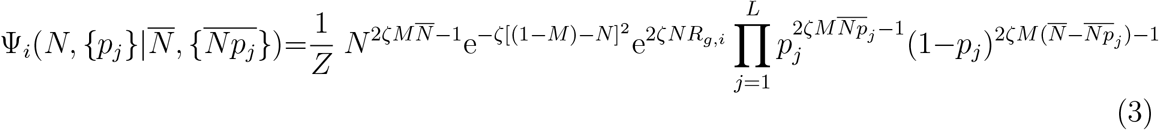

where *Z* is the normalisation constant.

### Numerical solution for the equilibrium

The state of each deme is determined by the average number of immigrant individuals 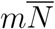 and of immigrant alleles 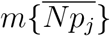 (at each locus *j*) per unit time. Given these, we can find the expected numbers and allel frequencies {𝔼_*i*_[*N*], 𝔼_*i*_[*Np*_*j*_]} on an island within habitat *i*, by integrating over the stationary distribution (eq. (3)). Here, the expectations 𝔼[] can be thought of as averages for a *given* island, obtained by averaging over replicate metapopulations (or simulations) or over uncorrelated timepoints at equilibrium within a single simulation. The crucial point is that at equilibrium, the average across all demes in the metapopulation at any instant (denoted by —) must equal the weighted sum of expected values across habitats (Rouhani and Barton, 1993). Thus:

**Table 1:**
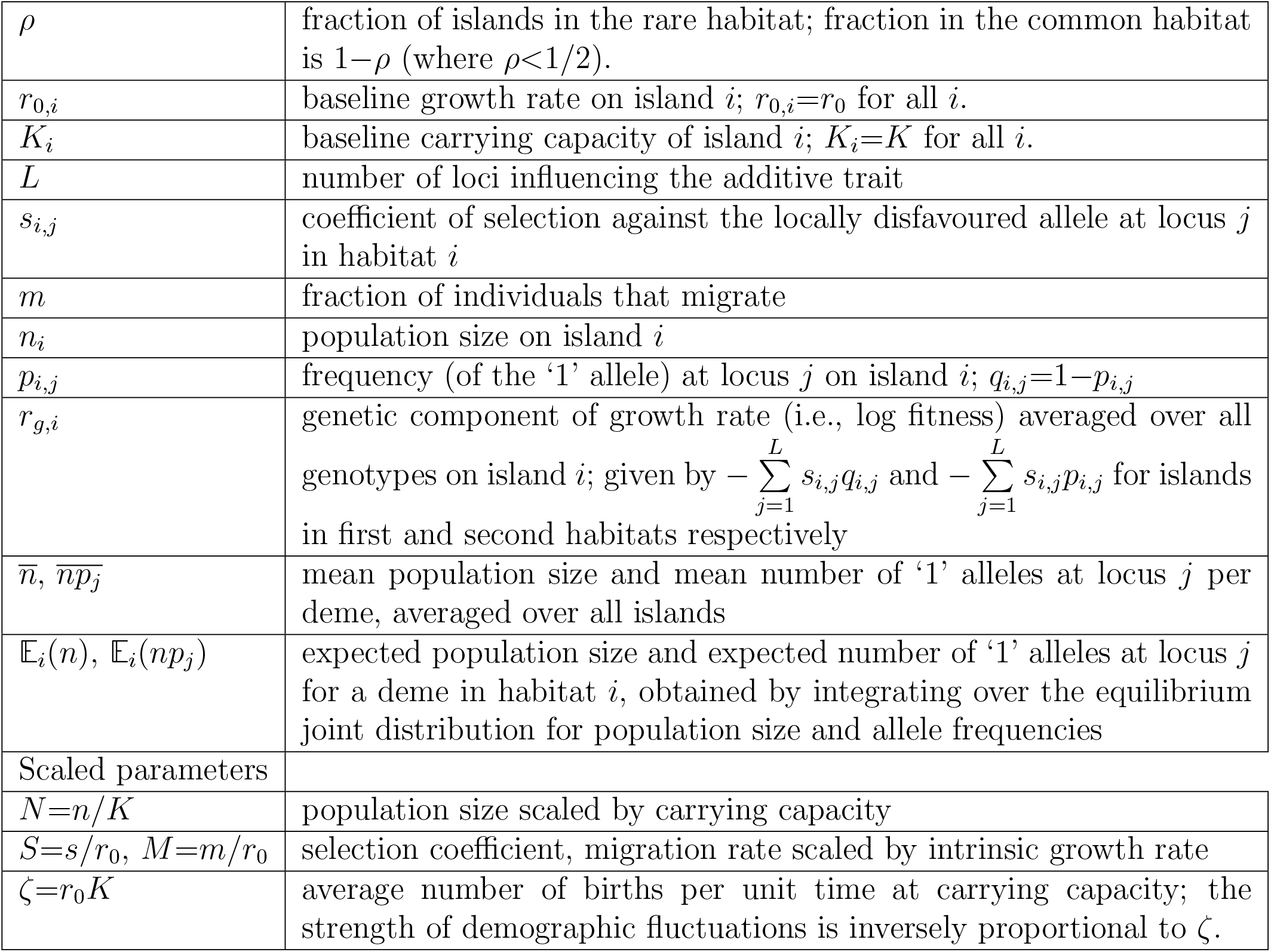
Key Notation

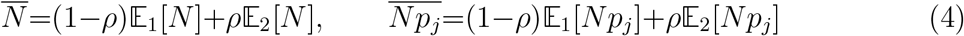

Equilibria are located by starting at an aebitrary 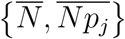, calculating {𝔼_*i*_[*N*], 𝔼_*i*_[*Np*_*j*_]} using eq. (3), then computing the new 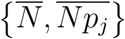 using eq. (4), and iterating until a fixed point. With this procedure, either a polymorphism is found, or one or other allele is fixed. In principle, this procedure simultaneously yields the equilibrium population size and allele frequencies at all the *L* loci (which may have different effect sizes and hence attain different frequencies). However, iterating over an *L*+1 dimensional space is computationally intensive. We thus restrict attention to the case with effect sizes equal at all loci, such that *S*_1,*j*_=*S*_1_ and *S*_2,*j*_=*S*_2_ for all *j*. Then, we need to find only the fixed point 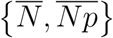.

In principle, populations may evolve towards different equilibria depending on their initial state. However, we find that outcomes are largely insensitive to initial frequencies (unless these are very extreme) except when migration rates are close to the threshold for loss of polymorphism and *LS* ≳1. Here, we will show the equilibrium that is attained starting with generalist populations (allele frequency 0.5 at each locus).

The above procedure for finding equilibria is exact, given the diffusion approximation which, however, relies on three assumptions. First, we assume all processes to be sufficiently slow (*r*_0_, *m, s* ≪ 1) that a continuous time approximation (eq. (1)) is valid. Second, we assume infinitely many demes, such that mean population size and allele frequency over the metapopulation exhibit negligible fluctuations, even though within any one deme, they follow a distribution (eq. (3)). This allows us to treat the migrant pool as deterministic, and completely characterised by 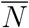 and 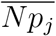. Finally, we assume that demes are in LE, i.e., LD between locally adaptive alleles at different loci is negligible. This final assumption is justified when recombination is much faster than other processes, i.e., as the unscaled parameters *s, m, r*_0_→ 0. We investigate the sensitivity of our results to each of these assumptions using individual-based simulations in the SI (Appendix C), but focus on the most critical assumption, namely that of LE, in the main text.

Since the full model involves several parameters, and calculating the joint distribution requires a numerical solution for 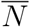 and 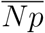, it is useful to consider some simpler limits. We first consider population dynamics in the absence of selection (*S*_1_=*S*_2_=0), and examine how demographic stochasticity and migration affect metapopulation survival. We then introduce selection, but assume it is weak relative to the baseline growth rate, i.e., *Ls*_*i*_ ≪ *r*_0_ or *LS*_*i*_ ≪ 1; we also neglect demographic stochasticity. In this ‘soft selection’ limit, population sizes are decoupled from fitnesses and both well-adapted and maladapted demes are at carrying capacity.

We then consider scenarios where selection against maladapted genotypes is strong enough to affect population dynamics, i.e., *LS*_1_, *LS*_2_ ∼1 (for equal-effect loci) resulting in ‘hard selection’, wherein less fit populations are smaller and may even go extinct.

We examine hard selection using the equilibrium distribution *ψ*(*N, p*_1_, … *p*_*L*_) derived above, as well as a simpler ‘semi-deterministic’ approximation which accounts for genetic drift but neglects demographic stochasticity, and treats population sizes as depending deterministically on mean fitnesses. In the main paper, we focus on symmetric selection across habitats (*S*_1_=*S*_2_=*S*); *S*_1_≠*S*_2_ is considered in SI, Appendix B.

## Results

### Effect of demographic stochasticity and migration in the absence of selection

Consider a scenario with no selection, such that population sizes are independent of allele frequencies, and only affected by demographic fluctuations and migration. Although individual demes fluctuate, 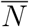 over the metapopulation evolves deterministically. Demes are coupled through 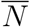, which determines the expected number of immigrants per deme. Using eq. (3), we can derive the distribution 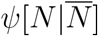 of population sizes *N*, conditioned on 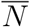 (SI, Appendix A3). The expected population size, 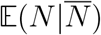 in any deme, given 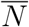, is obtained by integrating over 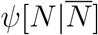, and then equating 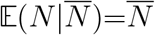. This yields one or more equilibria for 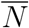.

There is always an equilibrium at 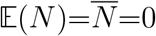, which corresponds to extinction of the whole metapopulation. Above a critical migration rate *M*_*crit*_, there may also be an equilibrium with 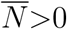. This *M*_*crit*_ is the migration rate at which the equilibrium 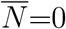 becomes unstable (SI, Appendix A3). Figure 1A shows that the *M*_*crit*_ required for metapopulation survival decreases exponentially: 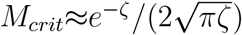 as *ζ* = *r*_0_*K* increases.

**Figure 1:**
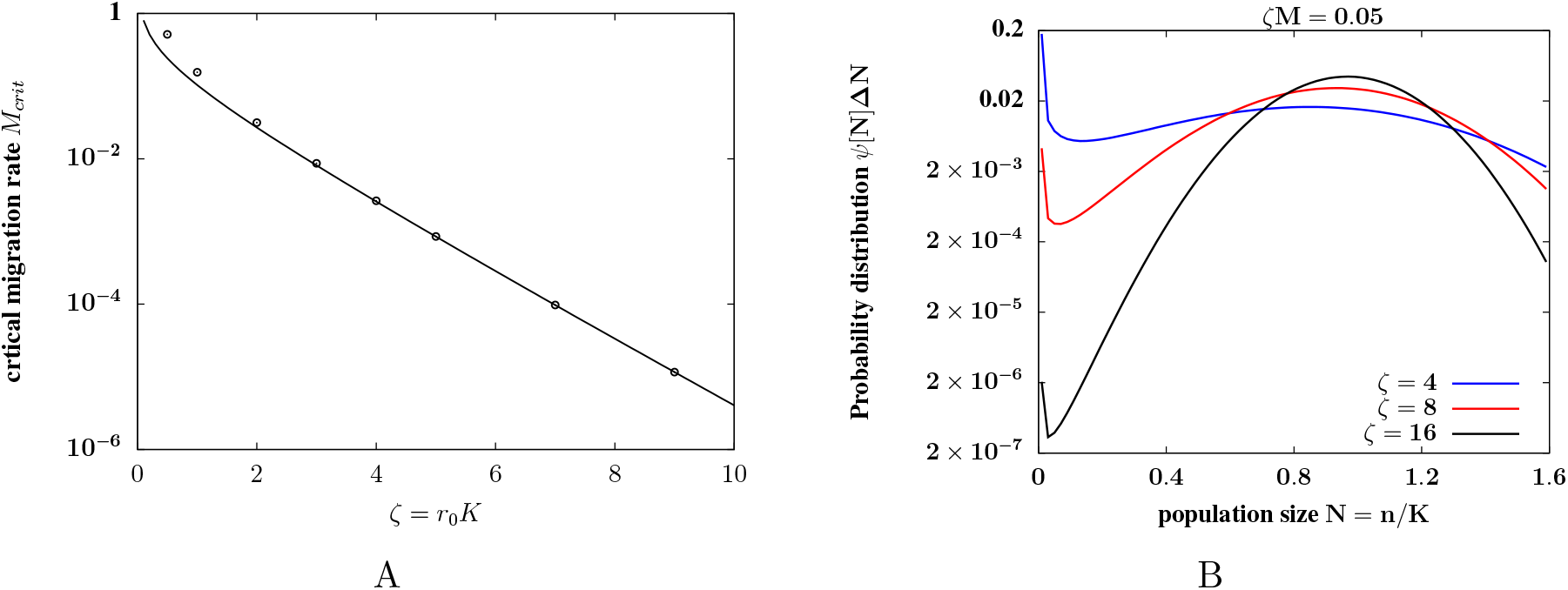
Deme sizes in the purely demographic model with no selection. (A) The migration threshold *M*_*crit*_, below which the entire metapopulation goes extinct due to demographic fluctuations, vs. *ζ*= *r*_0_*K*. The threshold *M*_*crit*_ falls exponentially with increasing *ζ*. Points show results of the diffusion approximation; the solid line depicts the large *ζ* approximation: 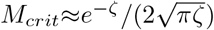. (B) Probability distribution *ψ*[*N*] of the scaled population size *N* =*n/K* (integrated over intervals of width Δ*N* =0.02), obtained from the diffusion approximation, for various *ζ*, for *ζM* =0.05. The migration rate *M* is reduced as *ζ* is increased, to keep 2*ζM* =*Km* fixed. The fraction of nearly extinct demes falls with increasing *ζ*. Diffusion approximation predictions are obtained from eq. S5 of the SI.

Above this migration threshold, the metapopulation survives as a whole; however, individual islands undergo extinctions and recolonisations if the average number of immigrants is small, i.e., for 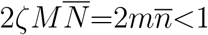, which corresponds to 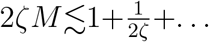 (SI, Appendix A3). In this case, the distribution of *N* is bimodal (fig. 1B): a fraction of demes is near extinction, whilst the remainder have population sizes normally distributed around *N* =1− *M* (i.e., *n*=*K*(1 − *m/r*_0_)), with variance 1*/*2*ζ*=1*/*(2*r*_0_*K*). The fraction of nearly extinct demes falls with increasing *ζ*, for a given 2*ζM* (fig. 1B). For migration rates that are still higher (2*ζM* »1), individual demes never go extinct and exhibit essentially deterministic dynamics.

The parameter *ζ*=*r*_0_*K* thus governs the extent of demographic stochasticity: it determines the threshold for global extinction (fig. 1A), the probability of local extinctions (fig. 1B), as well as the variance of population numbers around carrying capacity (among occupied demes). Henceforth, we consider growth rates and carrying capacities that are sufficiently high (i.e., *ζ*=*r*_0_*K* » 1) that local extinctions in the *absence* of maladaptation are exceedingly unlikely. However, even for *ζ*» 1, maladapted populations may be affected by demographic stochasticity and become extinct.

### Soft selection (constant population size)

We next introduce selection, but assume that the evolutionary change it effects is slow compared to population growth (i.e., *LS* ≪ 1), and that *ζ*=*r*_0_*K* » 1 (as above). Then the model reduces to the classical infinite island model with soft selection (Wright, 1932), where population sizes are fixed at carrying capacity (*n*=*K*) on each island. Unlike in the case with hard selection, allele frequencies at different loci evolve independently under soft selection (assuming LE), since genetic drift at any locus just depends on a fixed population size, and not on adaptation at other loci. Thus, we need only consider the allele frequency distribution *ψ*[*p*] at one locus: this was first derived by Wright (1932), and also emerges from the joint distribution in eq. (3) (SI, Appendix A4). This is also the basis of Blanquart et al.’s (2012) single-locus analysis.

The expected allele frequency 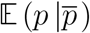 in a deme, given the mean 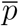 in the migrant pool, is obtained by integrating over *ψ*[*p*] (SI, Appendix A4). Allele frequencies in different demes are coupled via the mean allele frequency, 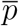, among migrants: within any deme, migration pulls the expected allele frequency towards 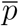, whereas selection drives 𝔼_1_[*p*] towards 1 (or 𝔼_2_[*p*] towards 0). Since all demes have equal sizes, they contribute equally to the migrant pool. Thus at equilibrium, we have 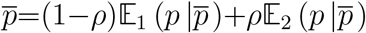, which allows us to numerically find the equilibrium allele frequency in each habitat and across the metapopulation (fig. 2A).

**Figure 2:**
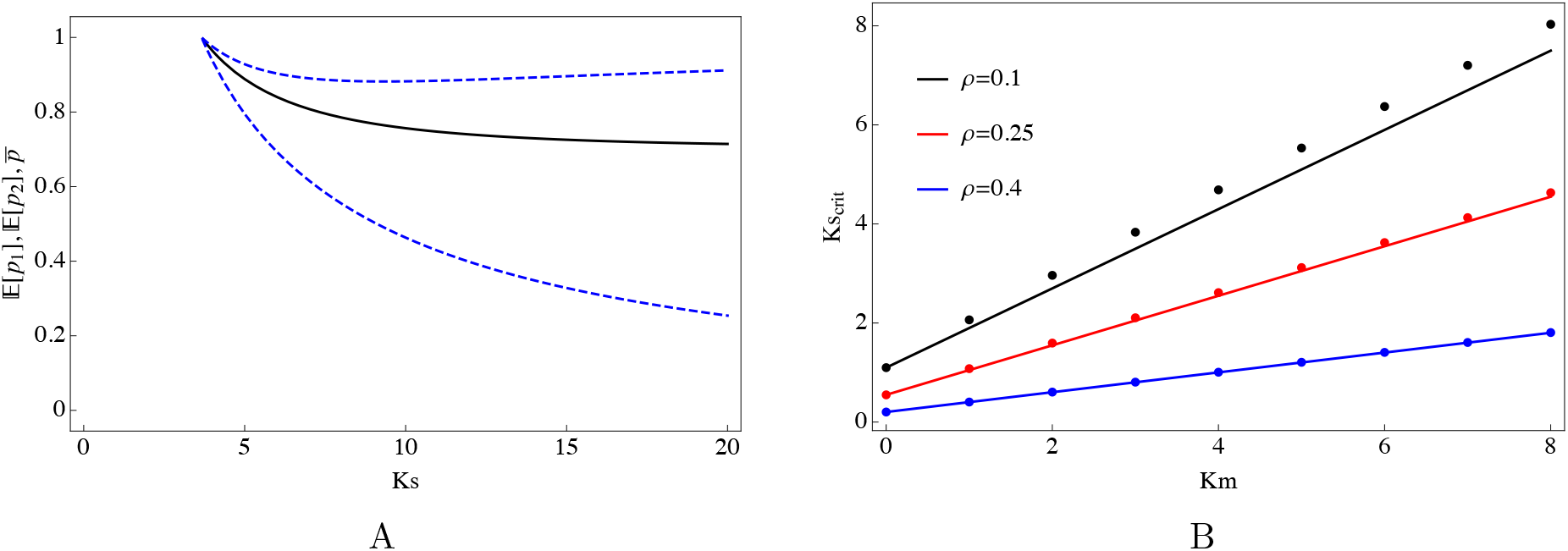
Local adaptation under soft selection. (A) Expected allele frequencies in the two habitats (dashed), and the overall mean over the metapopulation (solid), vs. *Ks*, as obtained from the diffusion. The fraction of demes in the rare habitat is *ρ*=0.3, the average number of migrants per generation is *Km*=8; selection is symmetric: *s*_1_=*s*_2_=*s*. (B) The critical selection coefficient *Ks*_*c*_, above which a polymorphic equilibrium with 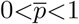 can be maintained, as a function of *Km*, the average number of migrants exchanged between demes, for various *ρ*. Points show the diffusion prediction; lines show the approximation 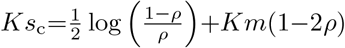. Allele frequencies under soft selection are obtained from the diffusion approximation by using eq. S6 in the SI.

There are always equilibria corresponding to 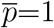 or 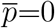 (i.e., when the whole metapopulation is fixed for one or other allele). A polymorphic equilibrium (with 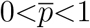) can be maintained when selection exceeds a threshold *s*_*c*_ (fig. 2A), such that alleles favoured in the rare habitat can invade. As selection becomes stronger, the different habitats approach fixation for different alleles. Thus, both habitats may be simultaneously adapted only if *s*>*s*_*c*_. For *s*<*s*_*c*_, alleles that confer a selective advantage in the common habitat fix across the entire metapopulation. Interestingly, the critical selection strength *s*_*c*_ approaches a non-zero value as *m*→ 0 (fig. 2B) as long as one habitat is rarer than the other. Thus, this bias towards alleles favoured in the common habitat persists even in the limit of very low migration, for which we would have (erroneously) expected allele frequencies in different demes to evolve independently.

To understand the dependence of *s*_*c*_ on *m*, it is useful to first consider a purely deterministic analysis, which ignores genetic drift (and thus requires *Ks, Km* » 1). This predicts that polymorphism can only be maintained above a critical selection coefficient *s*_*c*_=*m*(1−2*ρ*) (Appendix A4, SI), where *ρ* is the fraction of demes in the rare habitat. However, as *s* → *s*_*c*_, we expect 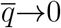 and thus 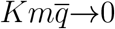 (fig. 2A). In other words, irrespective of how large *Km* is, near the threshold for loss of polymorphism, the numbers of alleles (of the rarer type) entering any deme will be very low and subject to random fluctuations due to drift. Thus the deterministic prediction only provides a lower bound on the true *s*_*c*_, as drift will further erode polymorphism.

In the opposite limit of low migration (*Km* →0), loci will be near fixation for one or other allele, making it necessary to account for drift. The rates of fixation towards and away from an allele with advantage *s*, at frequency 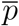 in the migrant pool, are in the ratio 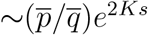, such that the expected frequency of the favoured allele in the deme is 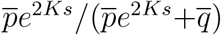 (SI, Appendix A4). Thus, with *s*_1_=*s*_2_=*s*, the metapopulation reaches an equilibrium at:

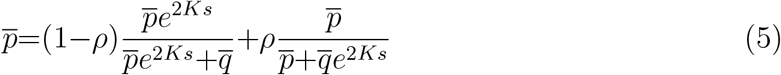

A polymorphic equilibrium at 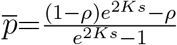 becomes possible if *Ks*>*Ks*_c_ where 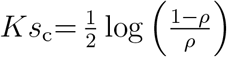 (in the *Km*→0 limit). Thus, even in the limit of very low migration, selection must exceed a critical threshold to prevent swamping of locally favoured alleles in the rare habitat. Note that this effect is not captured by the deterministic analysis above (which erroneously predicts *s*_*c*_→0 as *m*→0).

Numerically computing the equilibrium allele frequency, we find that the threshold *s*_c_ increases linearly with *m* (points in fig. 2B). This is approximated by: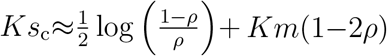 (solid lines in fig. 2B), where the first term is the *Km* → 0 prediction (which accounts for drift) and the second term the corresponding deterministic prediction (which neglects drift).

### Hard selection

When net selection against maladapted phenotypes is comparable to the baseline growth rate, i.e., *Ls* ∼*r*_0_ or *LS* ∼1 (assuming *L* equal-effect loci), changes in mean fitness have a substantial effect on population size, and maladapted populations are prone to extinction. Selection at *individual* loci need not be strong: typical effect sizes may be small enough (i.e., *ζS*=*Ks* ≤ 1) that drift can degrade adaptation at individual loci. However, if the number of loci *L* affecting fitness is large, selection, in aggregate, is strong. If local adaptation is to be possible even in large populations, then migration must be weak relative to selection per locus, i.e., *M* (1 − 2*ρ*)<*S*. Further, we assume *ζ*=*r*_0_*K* » 1, so that local extinction in the absence of maladaptation are highly improbable.

In the following, we use the joint distribution (eq. 3) to identify the conditions under which locally adapted, stable populations are maintained in both habitats. We first analyse two examples in detail to illustrate the qualitatively different outcomes under weak vs. strong coupling (feedback) between population size and mean fitness (fig. 3). We also compare theoretical predictions (which assume LE) with the results of individual-based simulations for these two examples, to clarify when LD can be neglected and the diffusion approximation becomes accurate (fig. 4). We then explore more generally how critical migration (or selection) thresholds for local adaptation depend on demographic stochasticity, the number of selected loci, and the fraction of the rare habitat.

**Figure 3:**
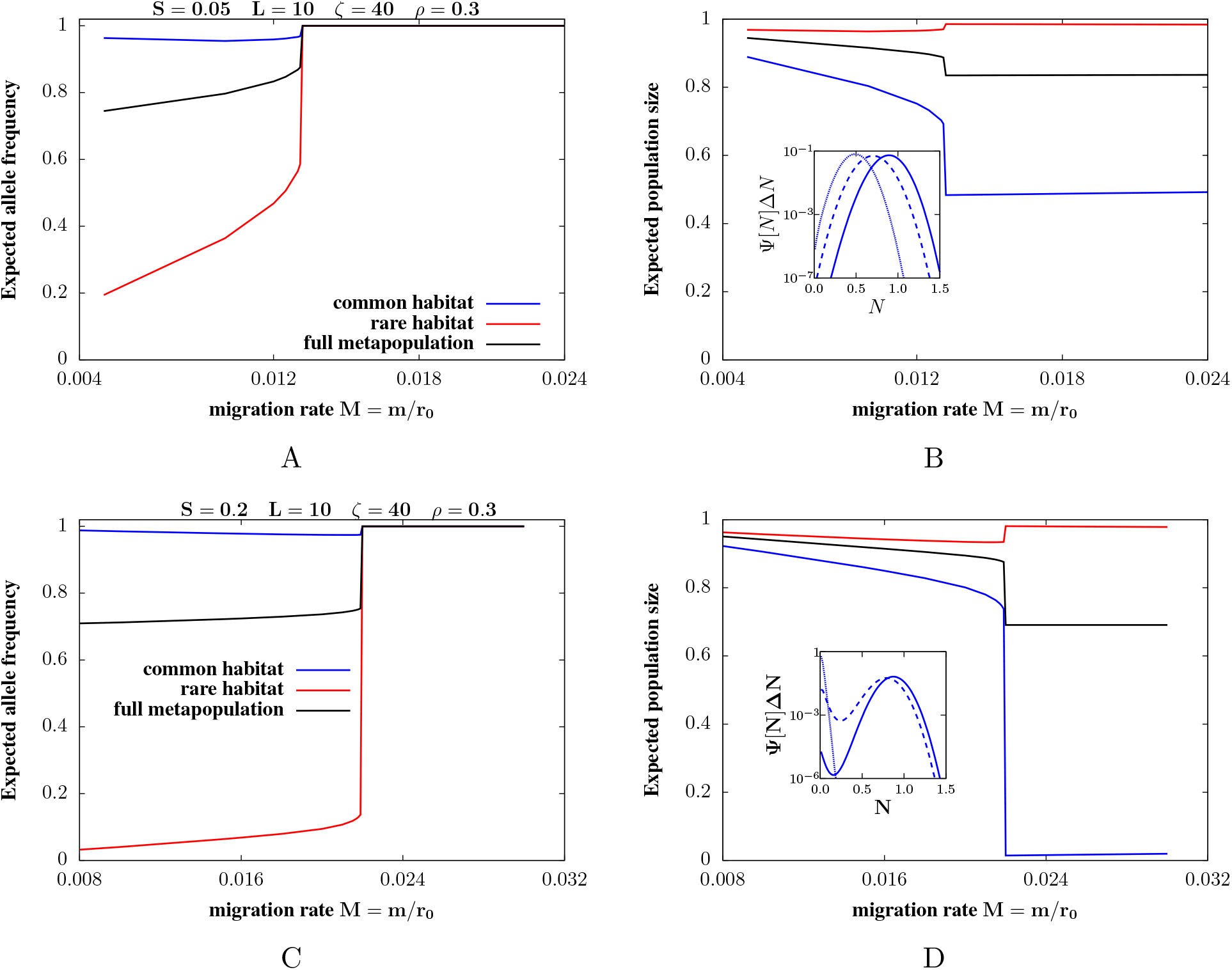
Loss of local adaptation at a critical migration rate under hard selection. Expected allele frequencies of the ‘1’ allele (left panels), which is favoured in the common habitat and disfavoured in the rare habitat, and expected population sizes (right panels) versus scaled migration rate *M* =*m/r*_0_, for (A)-(B) weak coupling (*S*=0.05, *LS*=0.5) and (C)-(D) strong coupling (*S*=0.2, *LS*=2) between population size and mean fitness. The number of selected loci is *L*=10 and selection is symmetric, with *S*_1_=*S*_2_=*S*=*s/r*_0_ at each locus; the rare habitat comprises 30% of demes (*ρ*=0.3) and *ζ*=*r*_0_*K*=40. The plots show the expected allele frequencies and sizes in the rare and common habitat (blue, red) as well as the mean 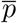 and 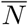 across the whole metapopulation (black). For both weak coupling (i.e., *LS*<1 in A,B) and strong (i.e., *LS*>1 in C,D), there is a critical migration threshold, *M*_*c*_, above which alleles favoured in the rare habitat are lost from the metapopulation. The insets in (B) and (D) depict the probability distribution *ψ*[*N*] of population sizes in the rare habitat (integrated over intervals of width Δ*N* =0.02) for *M*<*M*_*c*_ (solid line, corresponding to *M* =0.005 in B and *M* =0.014 in D), *M* ∼*M*_*c*_ (dashed line, corresponding to *M* =0.013 in B and *M* =0.0219 in D), and *M*>*M*_*c*_ (dotted line, corresponding to *M* =0.021 in B and *M* =0.03 in D). For weak coupling, *ψ*[*N*] peaks at a non-zero *N*, irrespective of *M*. For strong coupling, *ψ*[*N*] is bimodal for *M* ≲*M*_*c*_, with one peak close to *N* =0 (corresponding to nearly extinct demes) and the other peak at *N* ∼1−*LS*𝔼[*p*] (corresponding to a well-adapted demes). For *M*>*M*_*c*_, the second peak disappears and the distribution is concentrated at *N* =0 (all demes nearly extinct). All plots are obtained from the diffusion approximation by numerically determining fixed points (eqs. (3) and (4)) using the joint distribution Ψ[*N, p*].

**Figure 4:**
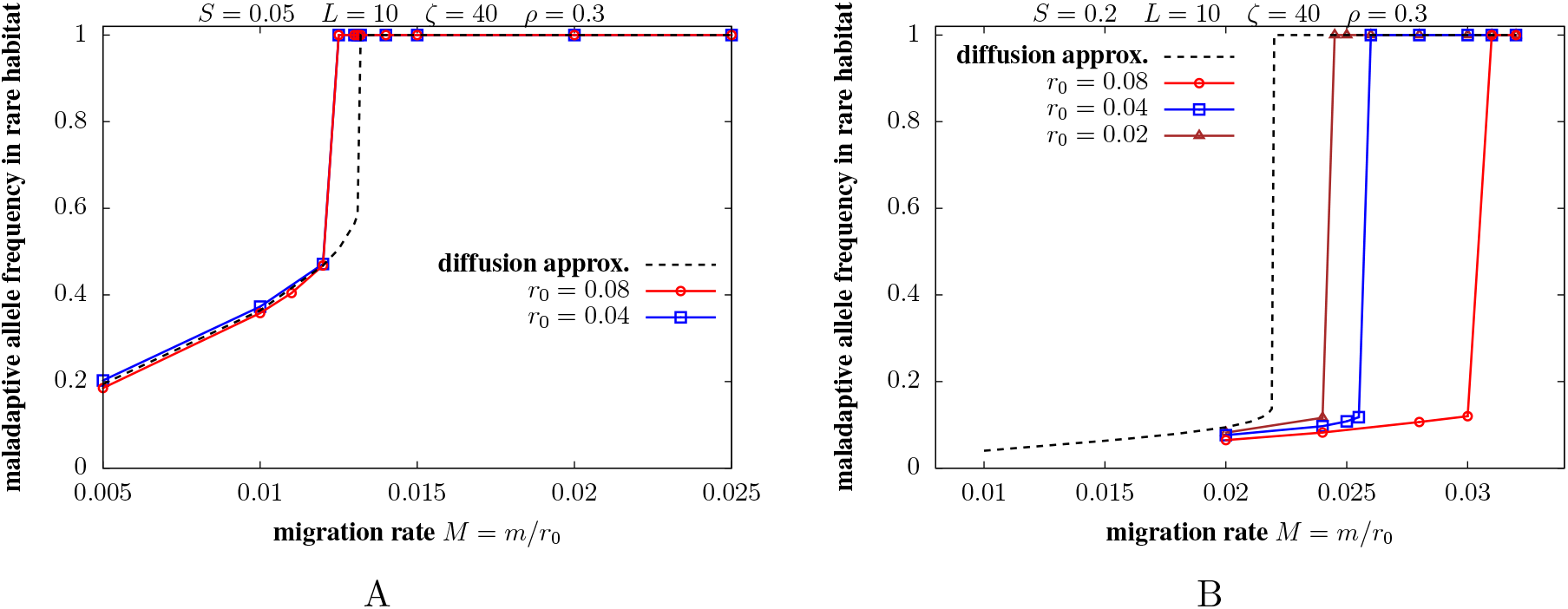
Comparison of the predictions of the diffusion approximation (which assumes LE) with individual-based simulations (which incorporate LD). Expected allele frequencies of the locally dis-favoured allele in the rare habitat versus scaled migration rate *M* =*m/r*_0_, for (A) weak coupling (*S*=0.05, *LS*=0.5) and (B) strong coupling (*S*=0.2, *LS*=2) between population size and mean fitness. All parameters are the same as in fig. 3. In each plot, the different colors show results of individual-based simulations (of 100 islands) for different values of *r*_0_. All other unscaled parameters *s, m, K* are varied (as described in the text) as we vary *r*_0_, such that the scaled parameters *S*=*s/r*_0_, *M* =*m/r*_0_ and *ζ*=*r*_0_*K* are the *same* for the different colors. The results of individual-based simulations deviate significantly from the diffusion prediction which assumes LE (dashed line) for larger *r*_0_, especially for *LS*>1 (panel B), but converge towards the LE prediction as we approach smaller *r*_0_ while holding scaled parameters fixed. We only simulate 2 values of *r*_0_ in panel A, as deviations from the LE prediction are already small even with the highest value of *r*_0_.

Figure 3 shows how polygenic adaptation collapses within the rare habitat as migration increases above a critical value in a scenario with weak coupling between population size and mean fitness, i.e., *LS*<1 (figs. 3A-3B) and in a strong coupling, i.e., *LS*>1 scenario (figs. 3C-3D). Figure 3 shows theoretical predictions for the expected allele frequencies and population sizes in the two habitats, as well as the distribution of population sizes in the rare habitat (insets). Figure 4 compares theoretical predictions with results of individual-based simulations.

In both the strong and weak coupling scenarios, alternative alleles are close to fixation in either habitat at low migration. As *M* increases, the frequency of the locally favoured allele (figs. 3A and 3C) and the expected population size *N* (figs. 3B and 3D) decline in both habitats due to increasing migration load. At a critical migration rate *M*_*c*_, the rarer allele is lost, the population in the rare habitat crashes, and the overall 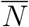 falls to a minimum. For *LS*>1, the loss of local adaptation results in near-extinction of the (maladapted) deme, while with weak coupling (*LS*<1), completely maladapted demes survive at a finite fraction of carrying capacity.

As *M* increases beyond *M*_*c*_, populations in the rare habitat increase marginally, signifying that this habitat is now a maladapted demographic sink. The emergence of source-sink dynamics at high *M* causes numbers in the common habitat to decline slightly with *M*. This is outweighed by the faster increase in numbers in the rare habitat, resulting in a slight increase in overall 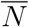 at large *M*. There is another migration threshold below which the whole metapopulation collapses because colonisation is too rare (fig. 1A); however, this threshold is negligibly small 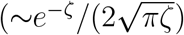 for large *ζ* (here, *ζ*=40), and is not visible here.

Figures 3B and 3D also depict how the distribution *ψ*[*N*] of the (scaled) population size in the rare habitat changes across the threshold *M*_*c*_ (insets). With weak coupling (*LS*< 1), population sizes are approximately normally distributed about a non-zero expected value 𝔼[*N*] irrespective of local adaptation, i.e., for both *M*<*M*_*c*_ and *M*>*M*_*c*_ (inset, fig. 3B). Further, 𝔼[*N*]~1− *LS*𝔼[*p*], where 𝔼[*p*] is the expected allele frequency of the locally deleterious allele in the rare habitat. By contrast, with strong coupling (i.e., *LS*>1), the distribution is bimodal for *M* ≤ *M*_*c*_ (i.e., when the rare habitat is locally adapted): a small fraction of demes is nearly extinct and the remaining have numbers that are approximately normally distributed around 𝔼[*N*]∼1−*LS*𝔼[*p*]. The fraction of nearly extinct demes in the rare habitat increases on approaching *M*_*c*_ (solid vs. dashed distribution, inset fig. 3D). Above the threshold *M*_*c*_, the distribution collapses to a single peak at *N* =0 (i.e., all demes are nearly extinct) and decays exponentially with *N*. The threshold for loss of local adaptation is sharper for larger *LS*— a finding that we clarify below.

The theoretical results shown in fig. 3 are based on the joint distribution of population size and allele frequencies (eq. (3)), derived by neglecting LD. As discussed above, we expect LD to be negligible and our analytical predictions to hold exactly in the limit where recombination is much faster than all other processes. This corresponds to taking the limit *s, m, r*_0_→0, *K*→∞, while holding the *scaled* parameters *S*=*s/r*_0_, *M* =*m/r*_0_ and *ζ*=*r*_0_*K* fixed.

To test this expectation, we compare theoretical predictions for the expected allele frequency in the rare habitat with individual-based simulations where the unscaled parameters *r*_0_, *s* and *m* are progressively reduced and the carrying capacity *K* increased, while holding fixed the scaled parameters *ζ*=*r*_0_*K, S*=*s/r*_0_ and *M* =*m/r*_0_ (note that the equilibrium is fully determined by scaled parameters under LE). These comparisons (figure 4) show that the migration threshold for loss of local adaptation in simulations can be significantly higher than the theoretical *M*_*c*_ if *LS* is high (fig. 4B). High values of *LS* correspond to high values of *Ls*, which governs the extent to which the *effective* migration rate of a deleterious allele is reduced due to its genetic background, when the two habitats are nearly fixed for alternative alleles at multiple loci (see also Appendix C, SI). However, as expected, the critical migration threshold in simulations converges to the theoretical prediction (for a given set of scaled parameters *ζ, S, M*) as evolutionary and ecological processes become weaker (*r*_0_, *s, m* → 0) relative to recombination. Note that for *LS*<1 (fig. 4A), the allele frequencies in individual-based simulations are very close to the theoretical (LE) prediction for all *r*_0_, suggesting that LD is already negligible even for the highest value of *r*_0_ (red). The minor discrepancy between simulations and theory in this case is due to the relatively few demes in the simulation, and vanishes as we simulate metapopulations with more demes (results not shown).

In the following, we will only present theoretical predictions obtained from equations (3) and (4) (which assume LE), with the understanding that these will accurately describe evolutionary outcomes in the well-defined limit in which LD is negligible.

### Semi-deterministic approximation

The fact that population sizes are approximately normally distributed about E(*N*) for *LS*≤ 1, suggests that in this ‘weak coupling’ regime, we can use a simpler approximation by neglecting fluctuations in *N* and assuming that, at any instant, it is close to its expected value 𝔼(*N*), which depends deterministically on the expected fitness 𝔼[*R*_*g*_] through 𝔼(*N*) ≈ 1+𝔼[*R*_*g*_]. This *semi-deterministic* approximation (details in SI, Appendix A5) thus accounts for how allele frequencies within any deme are influenced by genetic drift (whose strength is inversely proportional to the local population size *N*), but assumes that *N* itself is largely unaffected by demographic fluctuations and determined by the expected allele frequencies (through 𝔼[*R*_*g*_]). This approximation is thus only meaningful if *ζ* » 1, so that demographic fluctuations in *well-adapted* populations are weak. As shown below, the semi-deterministic approximation accurately predicts the threshold for loss of local adaptation if *LS* ≤1 (i.e., when the distribution of *N* is unimodal about the expected population size).

In this regime (i.e., when the semi-deterministic approximation is accurate), outcomes are governed by three parameters. For a given *ρ* (i.e., fraction of demes in the rare habitat), and assuming symmetric selection *S*_1_=*S*_2_=*S*, these are: *ζS*=*Ks*, which governs the strength of drift relative to selection in a population at carrying capacity, *ζM* = *Km*, which determines the average number of migrants exchanged between demes at carrying capacity, and *LS*, which determines how much population sizes are reduced below carrying capacity due to maladaptation (see also Appendix A5, SI). Below we clarify the roles of these parameters in the low migration limit *ζM* =*Km* ≪ 1, which is most conducive to local adaptation, and then build upon this to understand the more complex scenario where gene flow impairs adaptation more strongly.

### Low migration limit

Under rare migration, loci are near fixation for one or other allele within a deme. As with soft selection, this implies that the fixation rates of alternative alleles (at a given locus) on island *i* are in the ratio 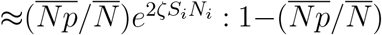, where *ζ*=*r*_0_*K*, and *S*_*i*_ is the (scaled) selective advantage of the locally favoured allele at that locus on island *i*. Further, 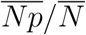 is the frequency (within the migrant pool) of alleles favoured on island *i*. A comparison of this heuristic (for fixation rates) under hard selection with the analogous approximation under soft selection (eq. (5)) highlights two important features of allele frequency evolution under hard selection.

First, the rate of fixation and hence the frequency of the favoured allele at any locus within any deme depends on the degree of maladaptation at all other loci via the local population size *N*_*i*_. In particular, locally deleterious alleles at very many loci, at even modest frequencies, can have substantial effects (in aggregate) on mean fitness, thus reducing size. This further accentuates drift at individual loci, causing locally deleterious alleles to increase or even fix, further reducing population size, thus generating a positive feedback between loss of fitness and decline in numbers.

Second, any island contributes to the allele frequency 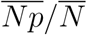 in the migrant pool in proportion to its size, which depends on the fitness of the island. Since locally adaptive alleles are at slightly lower frequency in the rare as opposed to the common habitat (even when both are locally adapted), demes are also somewhat smaller in the rare habitat. Thus, demes in the rare habitat contribute less to the allele frequency in the migrant pool than demes within the common habitat in proportion to the ratio of population sizes. This causes the allele frequency in the migrant pool to shift further towards the optimum for the common habitat, which increases migration load and reduces numbers in the rare habitat, generating a second positive feedback loop. Crucially, both kinds of feedback depend on the strength of coupling between population size and mean fitness, and are thus stronger for larger *LS*=*L*(*s/r*_0_).

Figure 5A shows how local adaptation depends on the the selective advantage of the favored allele per locus *S*=*s/r*_0_ or alternatively, the number of selected loci *L*, for a *fixed LS* under very weak migration (*ζM* =0.005). These plots thus reveal how local adaptation is influenced by the genetic architecture of (i.e., the number and selective effects of loci contributing to) genetic load, for a given total load *LS*, which translates into a given maximum possible reduction in population size under hard selection.

**Figure 5:**
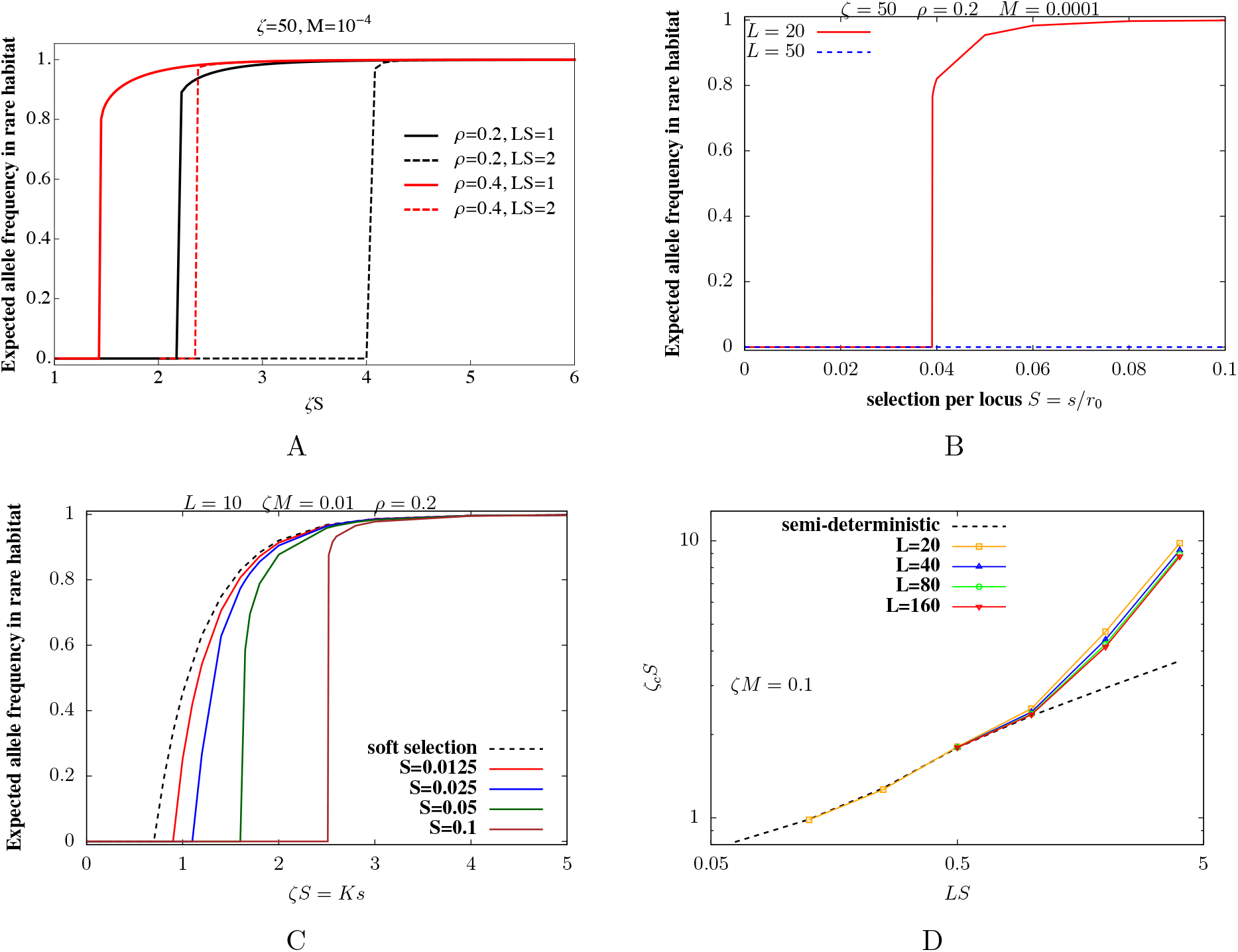
Local adaptation in the rare habitat under hard selection with weak migration (*ζM* =*Km*≪1); selection is symmetric across the two habitats (*S*_1_=*S*_2_=*S*=*s/r*_0_). (A)-(B) Expected frequency of the locally favoured allele in the rare habitat vs. *ζS*=*Ks*, for *ζ*=*r*_0_*K*=50, *M* =0.0001 for (A) fixed *LS* (B) fixed *L* (and *ρ*=0.2). (A) As we move along the x-axis, the scaled selection coefficient *S* and the number of loci *L* are changed simultaneously such that maximum possible genetic load *LS* is constant (for any one curve). Solid vs. dashed lines correspond to *LS*=1 and *LS*=2 respectively. Different colors correspond to different fractions *ρ* of demes in the rare habitat. Local adaptation in the rare habitat is lost when selective architectures are highly polygenic with weak selective effect per locus (high *L*, low *S*). (B) Local adaptation in the rare habitat is not possible at any selection strength *S*, for larger *L* (blue), which depends on *ζ*. (C) Expected frequency of the locally favoured allele in the rare habitat vs. *ζS*=*Ks* for different *S*=*s/r*_0_, for *ζM* =*Km*=0.01, *ρ*=0.2 and *L*=10. For a given *S*, the composite parameter *ζS*=*Ks* is varied by varying *ζ*; *M* is changed accordingly so that *ζM* is constant. The dashed line shows the corresponding prediction for allele frequencies as a function of *Ks* when population sizes are fixed, i.e., decoupled from fitness (soft selection). For a given *ζS*=*Ks*, populations approach the soft selection prediction as *S* and hence *LS* (which determines the coupling between population size and fitness) decrease. (D) Comparison of the predictions of the full stochastic model for the critical selection threshold *ζ*_*c*_*S* against the semi-deterministic prediction (which neglects demographic fluctuations and treats population size on each island as being determined by the local expected fitness). The threshold *ζ*_*c*_*S* for local adaptation in the rare habitat is plotted against *LS* for different values of *L* (depicted by different symbols), for *ζM* =0.1 and *ρ*=0.2. For given *L*, we vary *LS* by changing *S*, and then compute the critical *ζ*_*c*_ for each *S*. The migration rate *M* is changed accordingly such that *ζM* is constant at 0.1. The symbols and solid lines represent predictions of the full stochastic model while the dashed line represents predictions of the semi-deterministic approximation. There is good quantitative agreement between the full model and the semi-deterministic approximation for *LS*≲S1, but not for larger *LS*. Predictions of the full stochastic model with hard selection are obtained by determining fixed points numerically (eqs. (3) and (4)) using the joint distribution Ψ[*N, p*]; the soft selection prediction (dashed line in C) is obtained by determining fixed points under soft selection (eq. S6 in SI); the semi-deterministic prediction (dashed line in D) is obtained by determining fixed points of the semi-deterministic equations (eq. S7 in SI).

Figure 5A shows that given a certain maximum load *LS*, local adaptation in the rare habitat is possible only above a critical *S*_*c*_ per locus. For *S*<*S*_*c*_, drift overpowers selection at individual loci, causing alleles favoured in the common habitat to eventually fix across the entire metapopulation, despite very low genetic exchange, as with soft selection (fig. 2B). Alternatively, given a certain (maximum) load *LS*, local adaptation is possible only if the selected trait is determined by a modest number of loci (i.e., for *L*<*L*_*c*_, where *L*_*c*_=*L*(*S/S*_*c*_)), and fails for highly polygenic traits.

Further, local adaptation in the rare habitat requires stronger selection per locus when the total cost of maladaptation, *LS*, is higher (solid vs. dashed lines in fig. 5A). In fact, the critical selection threshold *S*_*c*_ increases with increasing number of loci under divergent selection, such that local adaptation not possible for *any S* for sufficiently large *L* even under weak migration (fig. 5B). As we argue below, higher *S* increases the efficacy of selection at individual loci (via *ζS*), but also results in stronger reduction in population *N* size due to load (via *LS*): the latter effect is especially strong for more polygenic traits, so that an increase in *S* may actually result in weaker selection relative to drift.

Thus, as selection becomes less ‘hard’, i.e., *LS*=*L*(*s/r*_0_) becomes smaller with *ζS*= *Ks* and *Km*=*ζM* held fixed, allele frequencies in the two habitats should approach those under soft selection. This is indeed what we see (fig. 5C): for a given *ζS*, the frequency of the locally favoured allele under hard selection *increases* towards the soft selection prediction on approaching lower *S* and consequently *LS*. Note that keeping *ζS* fixed corresponds to increasing carrying capacities as selection is reduced, such that the strength of selection relative to drift remains unchanged even as it becomes weaker relative to population growth.

Finally, *S*_*c*_ is lower for larger *ρ* (orange vs. blue plots in fig. 5A), i.e., if the rare habitat encompasses a larger fraction (but still less than half) of the islands. In this case, the rare habitat is subject to a lower migration load (since allele frequencies in the migrant pool are more intermediate), resulting in a weaker reduction in population size as well as weaker swamping at individual loci.

We now ask: for a given *ζM* =*Km*, does local adaptation depend only on the composite parameters *ζS*=*Ks* and *LS*, as one would expect under a semi-deterministic regime? To investigate this, we determine the threshold *ζ*_*c*_ (such that local adaptation occurs for *ζ*>*ζ*_*c*_), as a function of *S*, for various *L* and fixed *ζM*. Here, *ζM* is held constant by reducing the rescaled migration rate *M* =*m/r*_0_ as *ζ*=*r*_0_*K* increases, such that the average number of migrants (between demes at carrying capacity) remains unchanged.

Figure 5D shows that the semi-deterministic prediction for *ζ*_*c*_*S* (dashed line, obtained from eq. S7 of the SI) is extremely accurate for *LS*≲S1: in this regime, the threshold *ζ*_*c*_*S* for local adaptation in the rare habitat depends on the number of selected loci only via the combination *LS* (which governs the reduction in population size due to maladaptation). Moreover, this threshold only increases sub-linearly with *LS* for *LS*≳S1. By contrast, for *LS*)1, the semi-deterministic approximation fails: the critical *ζ*_*c*_*S* threshold increases much faster (nearly linearly) with *LS*, than predicted by the semi-deterministic approximation. However, even in this regime, the threshold for adaptation *ζ*_*c*_*S* depends only weakly on the number of loci, and is essentially governed by *LS*.

Note that the semi-deterministic approximation significantly under-predicts *ζ*_*c*_*S* (i.e., the extent to which selection per locus must prevail over drift) for large LS (fig. 5D). For *LS*>1, individual demes within the habitat may be maladapted and nearly extinct, even when the rare habitat is adapted as a whole. Thus, the distribution of population sizes is intrinsically bimodal (fig. 3D) and poorly approximated by a single population size (as assumed by the semi-deterministic approximation). Moreover demographic stochasticity may be important at low population numbers, even if it has negligible effects near carrying capacity (i.e., if *ζ*=*r*_0_*K* »1). Thus, accounting for the full *stochastic* distribution of population sizes and allele frequencies is crucial for predicting evolutionary outcomes in this regime.

### Critical migration rates for loss of local adaptation in the rare habitat

Where selection is strong relative to drift (for a given *LS*), so that both habitats are locally adapted under low genetic exchange, we ask: how high can migration be while still allowing adaptation to the rare habitat and polymorphism overall? Figure 6 shows *M*_*c*_, the critical migration rate above which polymorphism collapses, as a function of *S* for symmetric selection (*S*_1_=*S*_2_=*S*), for different values of *ζ* with *L* fixed (fig. 6A) or different values of *L* with *ζ* fixed (fig. 6B). The points represent results of the full stochastic model (based on the joint distribution of *N* and *p*); dotted lines show predictions of the semi-deterministic approximation (which neglects demographic stochasticity); the dashed line in fig. 6A shows the fully deterministic prediction (which neglects both demographic stochasticity and drift).

**Figure 6:**
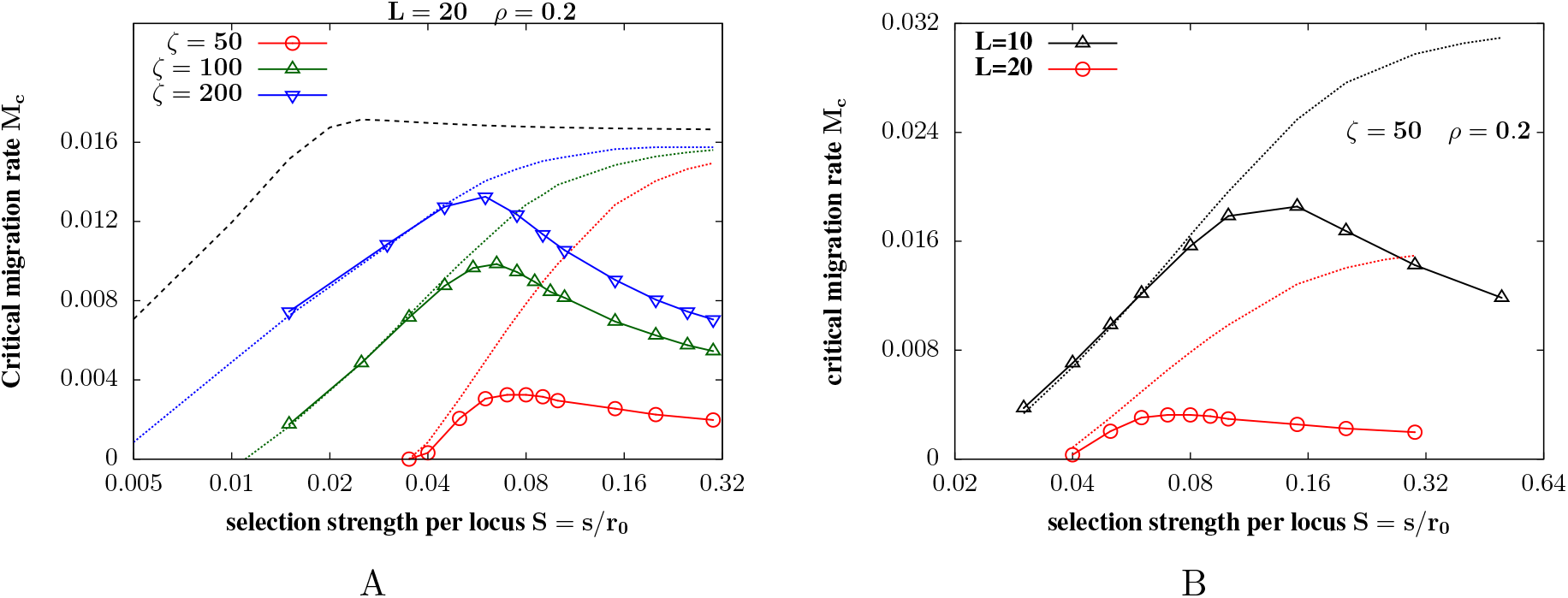
Critical migration rates for loss of local adaptation in the rare habitat. Critical (scaled) migration rate *M*_*c*_=*m*_*c*_*/r*_0_ versus (scaled) selection coefficient *S*=*s/r*_0_ per locus for (A) different values of *ζ* for *L*=20, (B) different values of *L* for *ζ*=50, with *S*_1_=*S*_2_=*S* and *ρ*=0.2 in both cases. Symbols with solid lines depict *M*_*c*_ obtained from the joint stochastic distribution of population size and allele frequencies; dotted lines represent the predictions of the semi-deterministic approximation (obtained from eq. S7 of the SI); dashed line in (B) represents the deterministic prediction. *M*_*c*_ falls with *S* for large *S*: this effect is not captured by the semi-deterministic or the deterministic prediction. For any *S*, the critical migration rate *M*_*c*_ increases with increasing *ζ* (in panel A).

It is useful to first consider the deterministic (dashed lines in fig. 6A) and semi-deterministic (dotted lines) predictions for the critical migration rate *M*_*c*_: both analyses predict that *M*_*c*_ should increase with *S* when selection is weak but then saturate to a constant value for large *S*. The semi-deterministic prediction for *M*_*c*_ is lower than the deterministic prediction due to the contribution of drift to maladaptation, but approaches the deterministic prediction as *ζ* (and hence *ζS*) increase. The emergence of a selection-independent threshold *M*_*c*_ at large *S* (under the deterministic analysis) is most easily demonstrated in the *ρ*→ 0 limit, in which one of the habitats is extremely rare and does not affect allele frequencies in the other habitat. In this simple limit, we can show that *M*_*c*_ ∼1*/*(4*L*) for *L* » 1 and large *S* (Appendix A6, SI). More generally, this reflects the fact that under hard selection, a population is viable only while its total migration load is less than its intrinsic growth rate *r*_0_. Since genetic load per locus is *at least m* in the limit of a very rare habitat (*ρ*→ 0), and is typically greater than *m* under hard selection (Appendix A6, SI), this limits how many polymorphic loci can be maintained without extinction.

Now, consider the predictions of the fully stochastic analysis (points in fig. 6A), which accounts for both drift and demographic stochasticity. When selection is weak, the critical migration rate *M*_*c*_ increases with *S* and is accurately predicted by the semi-deterministic analysis. However, beyond a certain threshold selection strength, which corresponds approximately to *LS*∼1 (fig. 6B), *M*_*c*_ *declines* with increasing *S*. Thus, the range of migration rates allowing local adaptation in the rare habitat is widest (i.e., *M*_*c*_ largest), for intermediate selection. As expected, *M*_*c*_ decreases with increasing *L* (in accordance with the difficulty of maintaining polygenic local adaptation under hard selection) and increases with increasing *ζ* (where higher *ζ*=*r*_0_*K* implies weaker stochastic fluctuations in both population sizes and allele frequencies). However, even for *ζ* as high as 200 (i.e., 200 births per generation in a well-adapted population), *M*_*c*_ is much lower than the corresponding deterministic threshold for large *LS* (fig. 6A).

The non-monotonic dependence of *M*_*c*_ on selection per locus appears quite generally, i.e., for various *ζ* (fig. 6A), *L* (fig. 6B) and *ρ* (results not shown), and is in sharp contrast to expectations under soft selection (where *M*_*c*_ is predicted to increase linearly with *S*), or to predictions for hard selection that fail to account for the stochastic distribution of population sizes (dashed and dotted lines in fig. 6). More specifically, loss of local adaptation in the large *LS* regime is accompanied by an increase in the fraction of nearly extinct demes; demographic stochasticity further exacerbates extinction risk in populations that may be already shrinking due to maladaptive gene flow, thus causing *M*_*c*_ to be several times lower than the deterministic prediction.

## Discussion

Existing theory on maladaptation and extinction in metapopulations is largely based on simple genetic models and fails to address the stochastic co-evolution of population sizes and allele frequencies. While various models include one or other stochastic process, e.g., genetic drift in subdivided populations (Blanquart et al., 2012; Whitlock and Barton, 1997), stochastic extinction-recolonisation dynamics in metapopulations (Hanski and Mononen, 2011), demographic and environmental fluctuations in the absence of selection (Mangel and Tier, 1993; Lande et al., 2003; Black and McKane, 2012), very few model the combined effects of different types of stochasticity (though see Chevin et al. (2017)). Our modeling framework is based on a diffusion approximation for the joint evolution of population sizes and allele frequencies under hard selection (Banglawala, 2010; Barton and Etheridge, 2018), which we extend here to a metapopulation with multiple ecological niches. It assumes a polygenic architecture for local adaptation, and accounts for both genetic drift and demographic stochasticity. It predicts the full stationary distribution of population sizes and allele frequencies in different habitats, thus clarifying the conditions under which local adaptation is maintained across habitats under divergent selection, or conversely, those that result in maladaptation and extinction in rare habitats. The underlying approximations (especially, LE) can be formally justified as *r*_0_ →0 (fig. 4). Thus, they may not be accurate in typical populations, where growth rates may be high. Similarly, fluctuations in population size are assumed to be only due to demographic stochasticity, and so may be greatly underestimated. Nevertheless, our modeling approach captures key processes involved in local adaptation, and our approximations apply over a broader range. We aim at understanding fundamental processes, rather than precise prediction.

We identify two distinct reasons why local adaptation fails within a rare habitat. First, if selection on locally favoured alleles is weak relative to drift, then alleles favoured in the common habitat tend to fix across the metapopulation, even when migration is extremely rare, i.e., *M*→ 0 (fig. 5). A somewhat paradoxical consequence of this result is that in the low *M* limit, loci that are under weak divergent selection across habitats are expected to show *weaker* differentiation than neutral loci, due to a net bias towards alleles favoured in the abundant habitat in the former case. In practice, we expect the time scale for loss of divergence at weakly selected loci to increase as *M* →0. Thus, local adaptation may be metastable and the loss of polymorphism extremely slow in this regime: this is also observed in individual-based simulations (results not shown).

The loss of local adaptation due to swamping from the abundant habitat, even under weak migration, is not predicted by deterministic arguments, and requires selection (per locus) to be weak relative to drift. This drift-dominated regime also emerges with soft selection, where we obtain an explicit expression for the critical selection threshold required for polymorphism (eq. (5)1). This threshold depends on the relative proportions of the two habitats: increasingly stringent selection per locus is required to maintain local adaptation in the rare habitat as it becomes more marginal (i.e., *ρ* decreases; fig. 2B). Conversely, when the two habitats are equally abundant (*ρ*=1*/*2), the critical selection threshold goes to zero as *M*→ 0 (see also Blanquart et al. (2012), who analyse this case with soft selection).

Local adaptation in the rare habitat can fail, even when selection per locus dominates over drift, if migration exceeds a critical threshold. Such thresholds emerge quite generally also with single loci under soft selection that are subject to maladaptive gene flow. In the present model (with hard selection and multiple selected loci), the total migration load sets a more severe constraint: it must be sufficiently low that the population can still grow. Since migration load scales with the number of loci under divergent selection, moderate maladaptation at many loci is sufficient to cause the population to crash. Declining population sizes further reduce the efficacy of selection at individual loci via increased drift, but also result in stronger swamping, generating a positive feedback that extinguishes populations in the rare habitat. This feedback sets an upper limit on the migration rate (given *L*), or alternatively, on the number of loci that can be divergently selected across the two habitats (given *M*), while still allowing local adaptation in both. Interestingly, we find that the critical level of maladaptive gene flow that populations can withstand actually decreases with increasing selection per locus, if the number of divergently selected loci is large, such that *LS*≳1 (fig. 6). The decrease in local adaptation with increasing intensity of selection, at a given *M*, was also noted by Ronce and Kirkpatrick (2001) in their deterministic analysis of two habitats under stabilizing selection. However, in the regime where maladaptation leads to extinction, i.e., for *LS*≳1, critical migration rates are poorly predicted by deterministic analyses (fig. 6A) which fail to account for the stochastic distribution of population sizes and the risk of extinction (of individual demes) within habitats that are adapted as a whole.

Our analysis thus highlights the difficulty of maintaining more than a few divergently selected alleles across habitats under hard selection, especially when one habitat is rarer. Selection per allele must be sufficiently strong to prevent swamping from the common habitat (even under extremely weak migration), but not so strong that the total fitness cost of maladaptation (due to migration load at multiple divergently selected loci) overwhelms population growth, triggering extinction. Thus, with increasing *L*, the conditions under which populations in the rare habitat can escape one or the other mode of failure become increasingly restrictive (figs. 5A, 5B, 6B).

Note that in our model, polygenic adaptation is difficult because of the rather extreme form of environmental heterogeneity, in which any selected allele has opposite effects on fitness in the two habitats. In an alternative model, where fitness depends on traits under stabilizing selection towards habitat-specific optima (as in Ronce and Kirkpatrick (2001)), migration load would be independent of the number of trait loci in the large *L* limit (Barton and Etheridge, 2018), and would thus not constrain the number of polymorphisms that can be maintained under hard selection. In this case, local adaptation is accomplished via small and transient allele frequency differences at multiple loci.

A key result is that local adaptation in the rare habitat becomes more difficult as selection becomes more ‘hard’ (fig. 5C): hard selection depresses population sizes, causing both drift and migration to become stronger relative to selection at individual loci. Thus hard selection and random drift can substantially increase the damage that gene flow may cause - as, for example, when farmed fish escape into wild populations (Glover et al., 2017).

We focus here on the case where locally adapted populations are demographically stable. However, the joint distribution derived in eq. 3 can be used to explore alternative regimes. For instance, we might consider a metapopulation with many very small demes and frequent extinction (i.e., *ζ*=*r*_0_*K*∼1). The whole metapopulation can still adapt to *global* selection pressures (if migration is sufficiently high), even when selection within each deme is weaker than local drift. Indeed, Wright (1932) argued that such a ‘shifting balance’ allows efficient search across alternative adaptive peaks (see Rouhani and Barton, 1993; Coyne et al., 1997). However, it would not be possible for populations to adapt to local variations in environment between demes in this regime.

The framework presented here is quite general, and applies directly to a wider range of cases, e.g., when the metapopulation encompasses more than two habitats or patches with heterogeneous carrying capacities and/or growth rates. While we have focused on local adaptation, the framework can be used to address other questions. For example, the model extends to include dominance, and so could be used to understand how heterosis and inbreeding depression influence local extinctions.

Our analysis neglects LD between locally adaptive alleles, which facilitates simultaneous local adaptation over a wider range of migration rates than predicted by the diffusion (fig. 4), because sets of introgressing alleles from differently adapted populations are eliminated together, thus reducing the effective rate of gene flow (Barton and Bengtsson, 1986). This effect is especially marked in the strong coupling regime (fig. 4B), where there is a strong positive feedback between a reduction in migration load (due to LD) and an increase in population size. The effects of LD can be incorporated, at least approximately, within the diffusion framework via the heuristic of effective migration rates; we defer analysis to future work. It may also be possible to estimate the extent of local adaptation, and the extent to which it reduces effective gene flow, by observing how divergence and LD vary along the genome (cf. Aeschbacher et al., 2017).

Local adaptation in a metapopulation may lead to parapatric speciation, despite gene flow: as populations diverge, selection against introgressing alleles increases, reducing effective migration, and allowing further divergence. A key issue here is whether a heterogeneous environment will lead to distinct clusters, separated by strong barriers to gene flow, which eventually become good biological species. This may depend on the distribution of available habitats. If these are broadly continuous, and select along multiple environmental dimensions, then there may be substantial local adaptation without clusters being apparent. However, with distinct environments, local adaptation may lead to strong isolation, as multiple divergent loci become coupled together (Barton and De Cara, 2009). The framework developed here may be used to investigate how the distribution of selective challenges influences whether populations evolve as generalists, adapting to a range of local environments, or split into distinct and well-isolated species.

## SUPPLEMENTARY INFORMATION

### A. Miscellaneous analytical derivations and results

#### A1. Derivation of the coupled stochastic equations for the joint evolution of allele frequencies and population size in any deme

For weak growth, selection and migration (i.e., *r*_0_, *s, m*≪1), we can use a continuous time approximation for allele frequency and population size dynamics. The size *n*_*i*_ and the allele frequency *p*_*i,j*_ at the *j*^*th*^ locus on the *i*^*th*^ island satisfy the following *coupled* equations (eq. 1 in the main text):

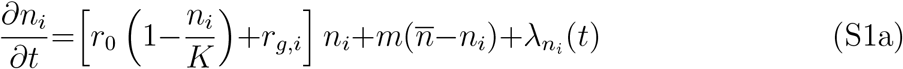

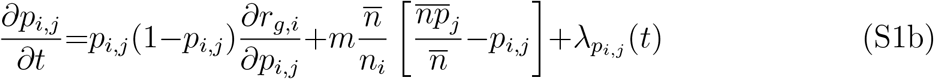

Here 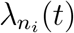 and 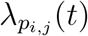 are uncorrelated random process with 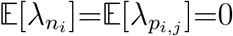 and 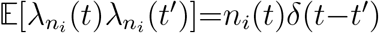, and 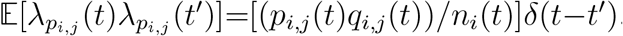.

Equation (S1a) describes population size dynamics on the *i*^*th*^ island in continuous time, and can be derived from the corresponding discrete-generation model, in which population size *n*_*i*_(*t*+1) in generation *t*+1 is Poisson distributed with mean 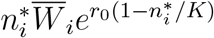, where 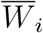 is the mean fitness on the island and 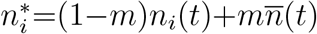 the population size just after migration.

The change in population size over one generation, *n*_*i*_(*t*+1)−*n*_*i*_(*t*), can be expressed as the *expected* change, 𝔼[*n*_*i*_(*t*+1)]−*n*_*i*_(*t*), plus a *noise* term Δ*n*_*i*_=*n*_*i*_(*t*+1)−𝔼[*n*_*i*_(*t*+1)]. The expected change is 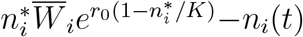. Taking the continuous time limit, which corresponds to *m, s, r*_0_→0, so that we need consider only lowest-order terms in these parameters, the expected change is: 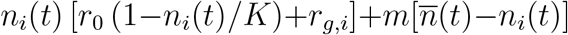. The properties of the noise term Δ*n*_*i*_ follow from the fact that population sizes are Poisson distributed in each generation. Thus, we have 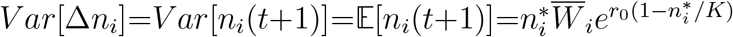, which is *n*_*i*_(*t*), to lowest order in *s, m, r*_0_. The white-noise property of the stochastic term in eq. (S1a) just reflects the fact that population size dynamics in the discrete generation model is Markovian, i.e., *n*_*i*_(*t*+1) depends only on *n*_*i*_(*t*).

The key simplification that results from taking the continuous time limit is that the change in population size (or allele frequency) under the combined influence of selection, migration, drift and demographic stochasticity can be approximated as the sum of the *individual* effects of each of these processes. Thus, the change in allele frequency at the *j*^*th*^ locus on the *i*^*th*^ island over time (eq. (S1b)), can be expressed as the sum of three terms, which reflect the change due to selection, migration, and drift.

The change due to selection at locus *j* (first term in eq. (S1b)) is the genic variance at that locus multiplied by *∂r*_*g*_*/∂p*_*j*_, which in the continuous time (*s*→0) limit, is just *s*_1,*j*_ and −*s*_2,*j*_ in the two habitats respectively. The change due to migration (second term) can be derived by expressing *p* at any locus as *n*_1_*/n* (where *n*_1_ is the number of individuals carrying the ‘1’ allele at that locus), then expressing Δ*p* as (Δ*n*_1_*/n*) − (*n*_1_*/n*^2^)Δ*n* (obtained via chain-rule differentiation), and finally using 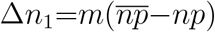 and 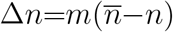. This yields the second (migration) term in eq. (S1b). The third term which describes stochastic fluctuations due to drift, and has a standard form that arises under Wright-Fisher sampling.

#### A2. Potential function for the joint distribution of population size and allele frequencies

Equation 2 in the main text specifies how the joint distribution Ψ_*i*_(**x**, *t*) of population sizes and allele frequencies (where **x** ≡ (*N, p*_1_, …, *p*_*L*_)) on any island in habitat *i* evolves through time. This equation is re-written below:

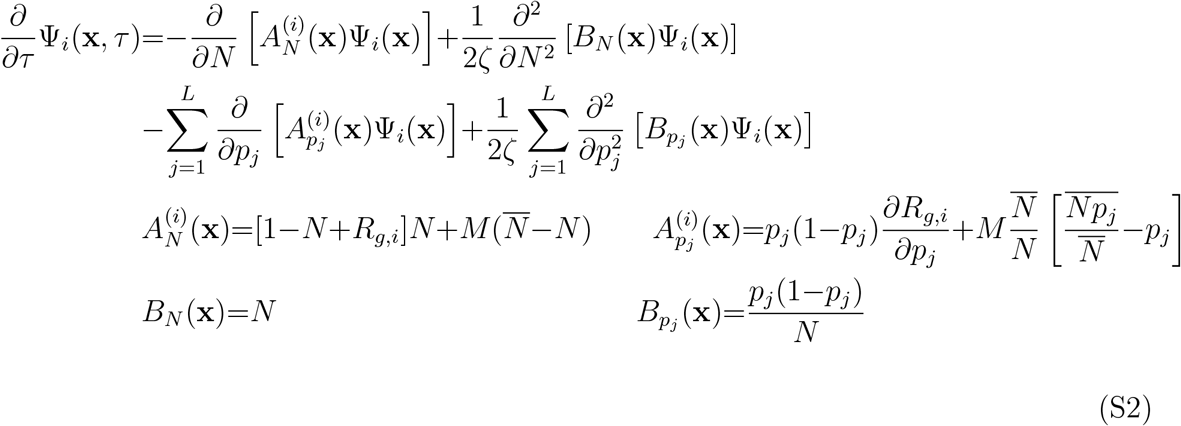

The equation is expressed in terms of the dimensionless parameters *τ* =*r*_0_*t, S*_*i,j*_=*s*_*i,j*_*/r*_0_, *M* =*m/r*_0_, and the composite parameter *ζ*=*r*_0_*K*, which is inversely proportional to the strength of stochastic fluctuations. It involves no mixed derivatives with respect to *N* and *p*_*j*_, as the covariance of fluctuations of *p* and *N* is zero (to first order in 1*/N, s*, etc.). The dependence on the habitat arises via the average log fitness *R*_*g,i*_=*r*_*g,i*_*/r*_0_, which is 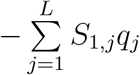 and 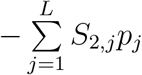 for *i*=1 (first) and *i*=2 (second) habitats respectively.

We can obtain an explicit solution for the *equilibrium* distribution in terms of a (suitably defined) potential function *U* (**x**):

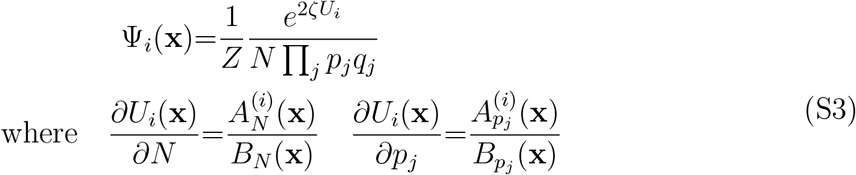

For such a solution to exist, we need 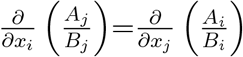, which requires density-independent selection and purely demographic stochasticity.

The potential function, obtained by integrating over *N* and {*p*_*j*_}, is given by:

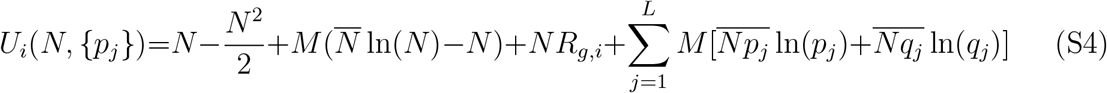

#### A3. Population size distribution with no selection

In the absence of selection (*S*_1,*j*_=*S*_2,*j*_ for all *j*), the joint distribution Ψ(*N*, {*p*_*j*_}) in equation 3 (main text) can be written as the product of independent distributions for *N* and each of the *p*_*j*_: allele frequencies at any locus follow Wright’s neutral distribution under the infinite island model (Wright, 1932). The distribution *ψ*[*N*] of deme sizes *N* is determined by 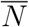, the mean population size across the entire metapopulation, and is given by:

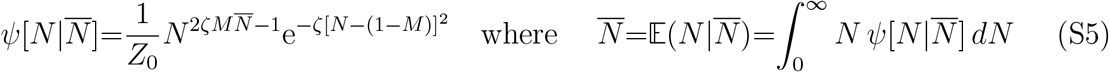

Here, *Z*_0_ is the normalisation constant. Integrating over 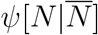 yields the expected population size 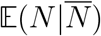 in any deme, given the mean 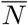; the equilibrium can be found by equating 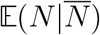 with 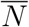.

We have checked this stationary distribution against both simulations of the discrete time model, and the prediction from the transition matrix. For example, for *r*_0_=0.05, *K*=100, *m*=0.01, the relative error in probability density is less than 3% in the bulk of the distribution, and the probability of extinction differs by 1.8%. Details are given in the accompanying Mathematica notebook.

There is always a solution 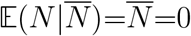, which corresponds to global extinction. Above a critical migration rate *M*_*crit*_, a second solution, corresponding to a non-zero mean population size emerges. The threshold *M*_*crit*_ is also that migration rate for which the solution 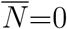 just becomes unstable, i.e., where 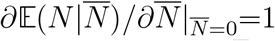. This yields the following equation for *M*_*crit*_: 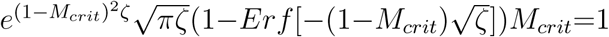. For large *ζ*, the threshold *M*_*crit*_ must be correspondingly small, such that we need only retain first order terms in*M*_*crit*_, and can further approximate 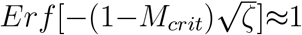. This yields 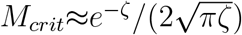 for large *ζ*. Only for *M>M*_*crit*_ can the metapopulation escape global extinction.

The shape of the distribution *ψ*[*N*] is governed by the parameter 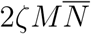, which is the average number of immigrants per generation per deme. For 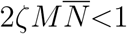, the distribution is bimodal: a fraction of demes support large populations with *N* ∼ 1 − *M*, while the remaining fraction is close to extinction. Conversely, for 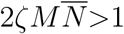, the distribution is unimodal: all demes support large populations, i.e., are close to carrying capacity.

We can obtain an explicit expression for the migration threshold *M*_*∗*_ for which 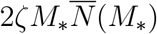 equals 1. Substituting the condition 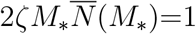 into the expression for 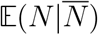 and then equating with 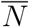 gives us an explicit expression for 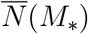. For large *ζ*, this is simply 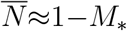. Substituting this into 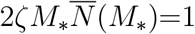, yields 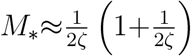. Thus, below this second threshold, i.e., for *M*_*crit*_*<M<M*_*∗*_, the stationary distribution of population sizes is bimodal, and there is a turnover of occupied vs. extinct demes due to frequent extinction and recolonisation, even at equilibrium. Note that even though there is always a non-zero fraction of nearly extinct demes for *M* such that *M*_*crit*_*<M<M*_*∗*_, this fraction declines rapidly with increasing *ζ* for a fixed 2*ζM* (fig. 1B in main text). Thus, even in this regime, one can practically ignore local extinctions for *ζ* ≫ 1 in the purely demographic model (i.e., with no selection).

#### A4. Thresholds for local adaptation under soft selection

In the limit *ζ* →∞, *S, M* →0 (with *ζS*=*Ks* and *ζM* =*Km* held constant), we recover the soft selection model, in which population sizes are decoupled from fitness and each deme is at carrying capacity (*N* =*n/K*=1), irrespective of adaptation. Allele frequencies now evolve independently of each other (assuming LE). In a deme where the favoured allele has selective advantage *s*, the allele frequency distribution is:

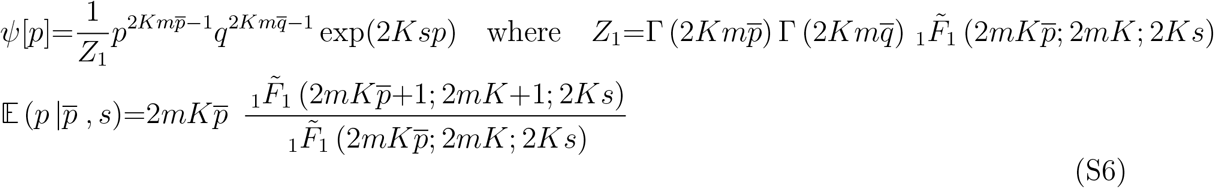

This distribution (Wright, 1932), can also be obtained by integrating the joint distribution in eq. 3 (main text) over *N*, and evaluating this integral as *ζ* →∞, *S, M* →0 for fixed *ζS*=*Ks* and *ζM* =*Km*. Note that the above distribution is expressed in terms of unscaled parameters *s, m* (since scaling to *r*_0_ is not meaningful under soft selection). One can now calculate the equilibrium frequency by equating the average allele frequency in the migrant pool 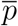 with the expected allele frequency (averaged over habitats) 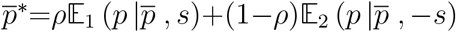, and solving numerically for the fixed point.

However, we can obtain approximate analytical solution in various limits. First, consider the deterministic limit (*Ks* ≫ 1 and *Km* ≫ 1), in which allele frequencies are tightly clustered about the expected value. Then the equilibrium can be found directly from eq. 1B (main text) by setting 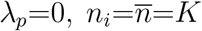 and 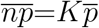. We can also obtain a prediction for *s*_*c*_, the critical selection required for a polymorphic equilibrium in the deterministic limit, by noting that just above the threshold *s*_*c*_ (i.e, when a polymorphic equilibrium first appears), the difference between allele frequencies in the two habitats must be very small, and the allele frequency in the common habitat close to 1. Thus retaining only lowest order terms in 1−*p*_1_ and *p*_1_−*p*_2_ in eq. 1B (main text) and solving for *p*_1_, *p*_2_ gives *s*_*c*_≈*m*(1−2*ρ*). Note that this prediction just provides a lower bound on *s*_*c*_, as we do not expect the deterministic analysis (which ignores variance of allele frequencies) to be accurate close to the threshold. Just above the threshold *s*_*c*_, we have 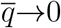, such that the distribution of allele frequencies is bimodal, making it necessary to account for drift. In general, we expect drift to further degrade local adaptation in the rare habitat, thus pushing the selection threshold *s*_*c*_ above the deterministic prediction. The opposite limit is that of weak migration *Km* → 0, in which any locus (within a deme) is nearly fixed for one or other allele. The rates of fixation towards and away from an allele with advantage *s*, which is at frequency 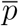 in the migrant pool, are in the ratio 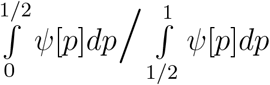 (for a deme in the rare habitat). Since most of the weight of the distribution *ψ*[*p*] is concentrated near *p*=0 and *p*=1 (for *Km* ≪ 1), we can approximate the integrand in the numerator (or in the denominator) by Taylor expanding *ψ*[*p*] near *p*=0 (or *p*=1). It then follows that the rates of fixation towards and away from the favoured allele are in the ratio 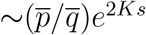. The expected frequency of the locally favoured allele is simply the (normalised) probability of fixation of this allele (eq. 7 in the main text).

#### A5. Semi-deterministic approximation for *LS*≲1

Under weak coupling (*LS*≲1), population sizes are approximately normally distributed about the expected value 𝔼 (*N*), irrespective of local adaptation (inset, fig. 3B, main text). This allows us to approximate the stationary distribution using a simpler ‘semi-deterministic’ approximation: we treat population size as being determined by the expected frequencies (i.e, neglect fluctuations in *N*), and assume that allele frequencies are distributed according to this deterministic population size.

In practice, we replace the population-averaged log fitness *r*_*g*_ by its expected value 𝔼[*r*_*g*_] in eq. 1A (main text), neglect the stochastic term *λ*_*n*_, and solve for the equilibrium size as a function of 𝔼[*r*_*g*_]. Further assuming that migration does not significantly affect population size, so that terms proportional to *m* in eq. 1A (main text) are also negligible, the scaled population sizes in the two habitats are 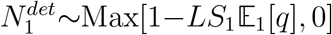 and 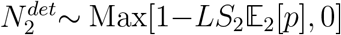: thus under the semi-deterministic approximation, population sizes are depressed below carrying capacity in proportion to the genetic load. To obtain 𝔼_1_[*p*] and 𝔼_2_[*p*], we assume that allele frequencies are distributed as in a model with fixed and deterministic population sizes 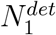 and 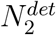. Then:

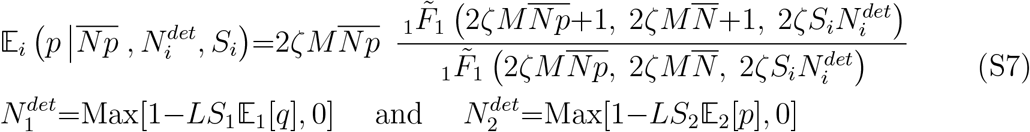

As before, the equilibrium 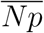 and consequently the expected allele frequencies can be determined iteratively from: 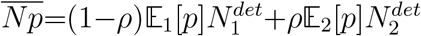.

#### A6. Deterministic analysis of the *ρ*→0 limit

To clarify the role of different kinds of stochasticity, it is useful to compare with the predictions of a deterministic analysis, which neglects both drift and demographic stochasticity. We can obtain the deterministic predictions by setting the noise terms *λ*_*n*_ and *λ*_*p*_ in eqs. (S1a) to zero, and then solving for the equilibrium population size and allele frequencies in each habitat.

Here we focus on the limit *ρ*→0, which is mathematically equivalent to continent-island migration, since the rare habitat (which occupies a vanishingly small fraction of demes) is too small to affect allele frequencies and population sizes in the common habitat. Then we have *N*_1_=1 and 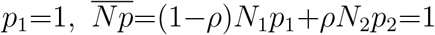 and 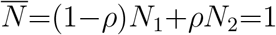. Substituting these into eqs. (S1), we have the following deterministic equations for the equilibrium allele frequencies and scaled population size in the rare habitat:

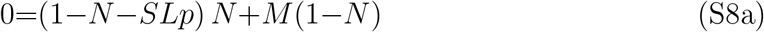

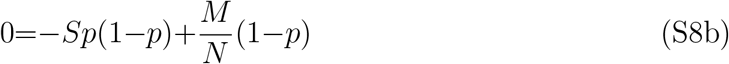

From eq. (S8b), we have *p*_*∗*_=*M/*(*NS*) for the polymorphic equilibrium in the rare habitat, which corresponds to a genetic load of *M/N* per polymorphic locus. Substituting into eq. (S8a), the equilibrium population size corresponding to the polymorphic equilibrium is: 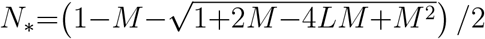. For the polymorphic equilibrium to be meaningful, we must have 0*<p*_*∗*_*<*1 and *N*_*∗*_*>*0. For *M/S<*1, this simply reduces to the requirement: 1+2*M* 4*LM* +*M* ^2^*>*0, which corresponds to *M<*1*/*4*L* for *L* ≫ 1. Thus the deterministic analysis predicts a critical migration threshold that is independent of *S* (for large *S*), and is inversely proportional to the number of loci under divergent selection.

### B. Asymmetric selection across habitats

In the main text, we detailed the symmetric selection scenario where alternative alleles are favoured in the two habitats, but selection per locally favoured allele is the same in each habitat (*S*_1_=*S*_2_). Here, we briefly consider the more general scenario where alternative alleles at a given locus may be selected more or less strongly in one of the two habitats (*S*_1_/=*S*_2_). For simplicity, we still assume selective effects to be equal across loci within a given habitat (i.e, *S*_1,*j*_=*S*_1_ and *S*_2,*j*_=*S*_2_ for all *j*).

Figure S1 depicts (*S*_1_, *S*_2_) combinations that allow simultaneous local adaptation (i.e., a polymorphic equilibrium) across the two habitats. For a given selection strength per locus, *S*_1_, in the common habitat, we determine two selection thresholds, *S*_2,*a*_ and *S*_2,*b*_, such that polymorphism is only possible for *S*_2_ (i.e., selection strengths per locus in the rare habitat) lying between these thresholds. For *S*_2_*<S*_2,*a*_, alleles favoured in the common habitat fix, while for *S*_2_*>S*_2,*b*_, alleles favoured in the rare habitat tend to fix. Note that the threshold *S*_2,*b*_ exists, i.e., strong selection in favour of alleles in the rare habitat can drive fixation of such alleles across the entire metapopulation, only if selection per locus in the common habitat is sufficiently weak.

Figure S1 shows the two selection thresholds for soft selection (black), and hard selection involving 10 (blue) or 20 (red) divergently selected loci. As before, soft selection is most conducive to simultaneous local adaptation. Under hard selection, the parameter combinations allowing for polymorphism become more restrictive with increasing *L*: this is consistent with the fact that larger *L* corresponds to stronger coupling between mean fitness and population size, which increases extinction probabilities in one or other habitat. This effect is exacerbated at higher migration rates (see fig. S1b): in this case, there is no (*S*_1_, *S*_2_) combination for which polymorphism is possible with *L*=20 loci (for *ζ*=50).

**Figure S1:**
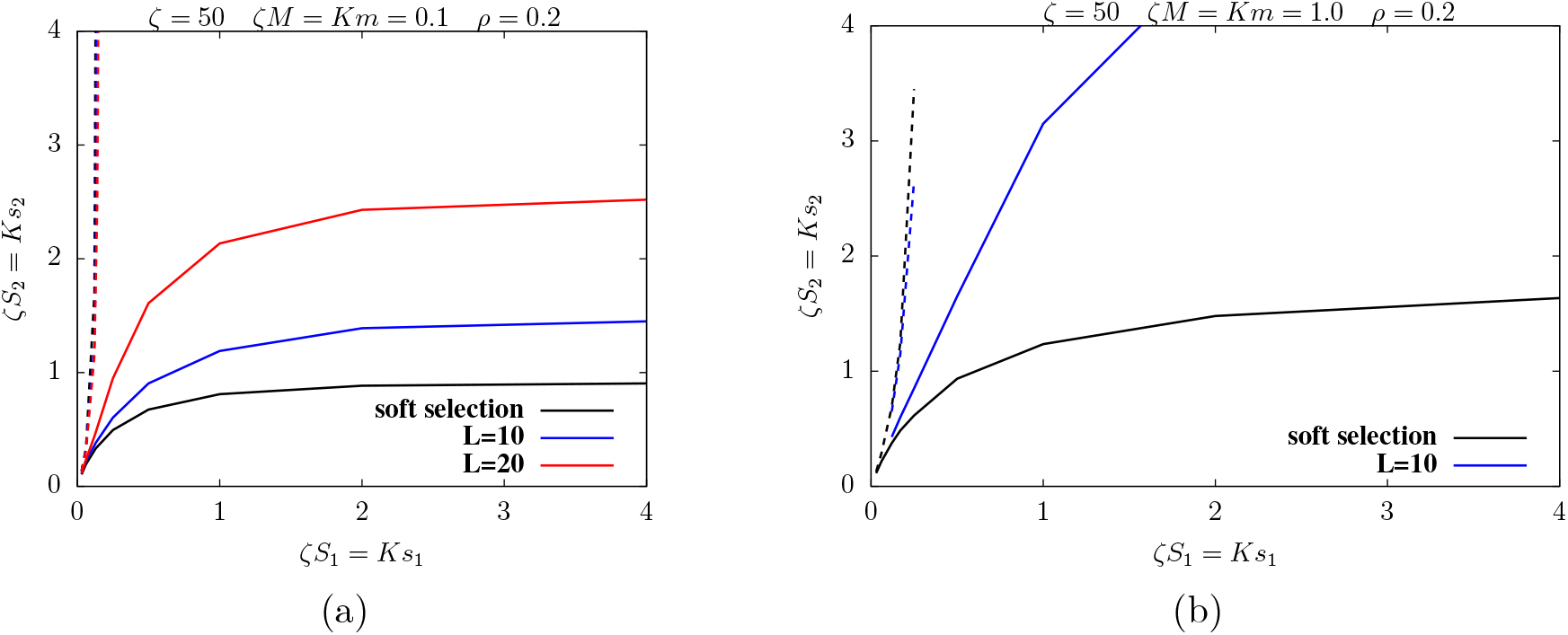
Local adaptation in scenarios with asymmetric selection across habitats. Critical selection thresholds *ζS*_2,*a*_ (solid lines) and *ζS*_2,*b*_ (dashed lines) in the rare habitat versus selection per locus, *ζS*_1_, in the common habitat for (A) low migration (*ζM* =0.1) and (B) intermediate migration (*ζM* =1.0). Alleles favoured in the common habitat fix across the entire metapopulation for *S*_2_*<S*_2,*a*_ (below solid line); alleles favoured in the rare habitat fix across the entire metapopulation for *S*_2_*>S*_2,*b*_ (above dashed line); a polymorphic equilibrium corresponding to local adaptation across the two habitats is possible for *S*_2,*a*_*<S*_2_*<S*_2,*b*_, i.e., for parameter combinations lying between the solid and dashed lines. The different colors correspond to different degrees of coupling between population size and mean fitness: soft selection, i.e., constant population sizes (black), hard selection with *L*=10 (blue) and *L*=20 (red) loci; larger *L* corresponds to stronger coupling (for fixed *ζS*_1_). The region in parameter space allowing for simultaneous local adaptation shrinks with increasing *L*; at sufficiently high migration, there is no (*S*_1_, *S*_2_) combination for which polymorphism is possible with *L*=20 loci in (B). Selection thresholds under hard selection are obtained by determining fixed points numerically (eqs. 3,4 in main text) using the joint distribution Ψ[*N, p*]; selection thresholds under soft selection are obtained by determining fixed points using the allele frequency distribution *ψ*[*p*] (eq. (S6)).

### C. Individual-based simulations

Our theoretical framework is based on the diffusion approximation, which involves three assumptions, namely, continuous time, an infinite number of demes and linkage equilibrium (see also Model and Methods). We test the validity of each assumption by comparing analytical predictions with two kinds of individual-based simulations.

In simulations of the *first* kind, we only simulate a single focal deme (belonging to the rare habitat) that is subject to immigration from the rest of the metapopulation, whose state is assumed to be fixed and accurately described by the diffusion framework. Migration into the focal deme is simulated by drawing the number of immigrants per generation from a Poisson distribution with mean 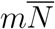, and assigning the ‘1’ allele at (any) locus *j* in each migrant genome independently with probability 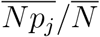, and ‘0’ otherwise. Here, 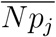 and 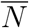 are the means across the whole metapopulation, and are obtained numerically from the diffusion approximation (as described above). These simulations respect the infinite island assumption, since migrants are drawn from an effectively infinite pool, characterised by deterministic 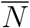 and 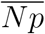. This set of simulations thus allows us to test the consequences of discrete time and LD within demes arising from the fact that allele frequencies in the focal deme differ from the average 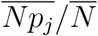 across the metapopulation at multiple loci. Note, however, that this is not an accurate representation of LD in the metapopulation, as it neglects structure within the migrant pool itself: this consists of two sets of genotypes (originating from the two habitats) that are differentiated from each other at multiple loci.

In simulations of the *second* kind, we simulate the full metapopulation consisting of *n*_*D*_ demes, of which (1 − *ρ*)*n*_*D*_ and *ρn*_*D*_ belong to the first and second habitat respectively. In these, we explicitly simulate full multi-locus genotypes of all individuals across all demes. Thus, the second set of simulations allows us to also examine the consequences of the infinite deme assumption (by varying *n*_*D*_) as well as the full effects of LD. Note that simulations of the second kind do not make any simplifying assumptions whatsoever, and explicitly account for the fact that the migrant pool itself consists of two groups of diverged genotypes when the two habitats are simultaneously adapted at multiple loci. Figure 4 in the main text shows results of individual-based simulations of the second kind.

Since individual-based simulations of large numbers of demes (e.g., in simulations of the second kind) with large carrying capacities (*ζ* »1) and polygenic architectures are computationally intensive, we focus on one set of parameters and explore how the critical migration threshold for loss of local adaptation is affected by deviations from these three assumptions for this set.

#### Continuous time approximation

To test the validity of the continuous time approximation, we simulate a single focal deme in the rare habitat subject to immigration from a migrant pool whose state is assumed to follow the predictions of the diffusion framework (simulations of the first kind). Figure S2a and S2b show the average allele frequency (averaged across all loci) and the average population size (averaged over 100 replicates) from individual-based simulations for various *r*_0_. The unscaled parameters *s* and *m* are decreased and the carrying capacity *K* increased as we reduce *r*_0_, such that the scaled parameters *ζ*=*r*_0_*K, S*=*s/r*_0_ and *M* =*m/r*_0_ remain constant. Figure S2a and S2b show that even for *r*_0_ ∼0.5, individual-based simulations only deviate moderately from the diffusion prediction (solid line), with the agreement between the two improving for lower *r*_0_.

The slightly higher frequencies of locally adaptive alleles in simulations (than predicted theoretically) at larger *r*_0_ may reflect the importance of 𝒪(*s*^2^) terms (which the continuous time approximation neglects) but also weak LD generated by migration, if 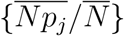 are significantly different from allele frequencies in the focal deme.

#### Finite number of demes

We also carry out individual-based simulations of metapopulations with a finite number *n*_*D*_ of demes (simulations of the second kind). Figures S2c, S2d) show simulations with 100 versus 500 demes (circles versus triangles) of which 20% belong to the rare habitat, along with theoretical (diffusion) predictions. For any given set of parameters, the average frequency of locally favoured alleles and average population sized in the rare habitat are slightly higher for *n*_*D*_=500 than for *n*_*D*_=100. Moreover, the migration threshold *M*_*c*_ itself is also somewhat higher for larger *n*_*D*_. However, these effects are modest in magnitude.

**Figure S2:**
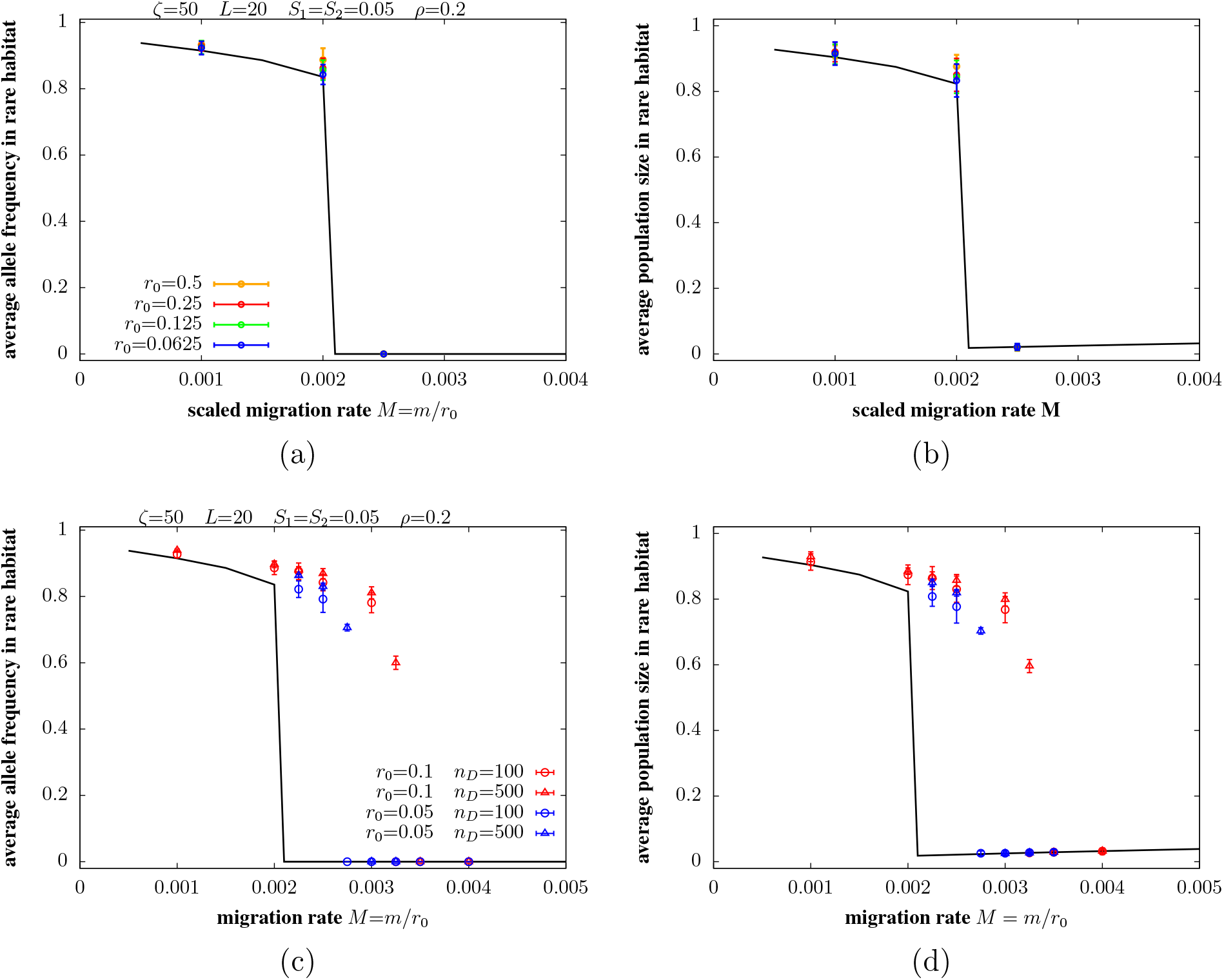
Comparison of individual-based simulations with predictions of the diffusion approximation (eqs. 3 and 4 in the main text). (A)-(B) Average allele frequency and population size in the rare habitat versus scaled migration rate *M* =*m/r*_0_ from individual-based simulations of a focal deme which receives a Poisson-distributed number of immigrants (on average 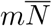) in each generation. Migrants are sampled from an infinite pool in LE; allele frequencies in this pool are equal to 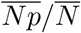 (at each locus). The quantities 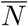 and 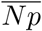 are calculated using the diffusion approximation for the infinite island model (eqs. 3,4 in the main text). Results of individual-based simulations are shown by circles, the different colors corresponding to different values of *r*_0_. All other unscaled parameters are varied as we vary *r*_0_ such that the scaled parameters remain constant at *ζ*=50, *S*_1_=*S*_2_=0.05. Solid line shows predictions of the diffusion approximation. (C)-(D) Average allele frequency and population size in the rare habitat versus scaled migration rate *M* =*m/r*_0_ from individual-based simulations of a metapopulation with *n*_*D*_ demes, where circles and triangles represent results for *n*_*D*_=100 and *n*_*D*_=500. For each *n*_*D*_ and *M*, we simulate two sets of unscaled parameters, one with *r*_0_=0.1 (red) and the other with *r*_0_=0.05 (blue), both corresponding to the same scaled parameters: *ζ*=50 and *S*_1_=*S*_2_=0.05. The solid black line shows predictions of the diffusion approximation (assuming LE and infinitely many islands). The frequency of the rare habitat is *ρ*=0.2 and the number of selected loci is *L*=20 in all simulations.

#### Linkage equilibrium

For a given *n*_*D*_, we further compare populations with different values of the baseline growth rate: *r*_0_=0.05, *K*=1000 vs. *r*_0_=0.1, *K*=500, (red vs. blue symbols), both of which are described by the same scaled parameters (*ζ*=50, *S*_1_=*s*_1_*/r*_0_= *s*_2_*/r*_0_=*S*_2_=0.05). As also observed in Figure 4B in the main text (for a different set of parameters), *M*_*c*_ is higher than the diffusion prediction (solid black line) for both sets of parameters, but appears to be converging towards the diffusion prediction as *r*_0_ → 0 (i.e., as all other processes become weaker than recombination).

As discussed in the main text and above, the diffusion approximation assumes that selection at any locus is independent of the allelic state at other loci, i.e., LD breaks down much faster than allele frequencies change. The fact that allele frequencies within a deme differ from *average* allele frequencies within the migrant pool at multiple loci appears to be of little consequence (figs. S2a and S2b). What matters is that the migrant pool consists of two very different sets of genotypes: thus, immigrant genotypes entering a particular deme may either be perfectly adapted (if they originate from other demes belonging to the same habitat) or severely unfit (if they emerge from demes belonging to the other habitat). Since deleterious immigrant alleles typically originate from the alternative habitat, they are embedded in highly unfit genetic backgrounds and will be rapidly eliminated, causing their effective migration rate to be much lower than *m*. This reduction in the effective migration rate of alleles that originate from the ‘wrong’ habitat, if significant, allows polymorphisms to be maintained over a much wider range of migration rates than predicted by the LE analysis (figs. S2c and S2d), especially in rarer habitats.

We can obtain an estimate for how much effective migration rates are reduced as follows: if populations in both habitats are nearly perfectly adapted, then an allele immigrating into a deme from the alternative habitat is associated with *L*− 1 other deleterious alleles and suffers a first-generation selective disadvantage that is ∼*e*^−2*s*(*L*−1)^. On the other hand, an allele immigrating from a deme in the same habitat suffers no such “extra” selective disadvantage. Thus, the long-term reproductive contribution of immigrant alleles depends not only on their own selective effect, but also on their habitat of origin. The reduction in reproductive value of alleles from the ‘wrong’ habitat (relative to alleles from the same habitat), given by *e*^−2*s*(*L*−1)^, becomes less pronounced as *sL* approaches zero. This limit is approached as we consider smaller and smaller unscaled parameters *r*_0_, *s*, … (red vs. blue symbols in figs. S2c and S2d), while holding the scaled selective effect *S*=*s/r*_0_ *constant*. In effect, in this limit, the associations between deleterious alleles are broken up sufficiently fast that we can assume LE over the time scales over which deleterious alleles are purged by selection. We defer an analysis of how the effects of LD can be incorporated within the diffusion framework to future work.

## Notes

### Competing Interest Statement

The authors have declared no competing interest.

## References

Aeschbacher, S., J. P. Selby, J. H. Willis, and G. Coop. 2017. Population-genomic in-ference of the strength and timing of selection against gene flow. Proceedings of the National Academy of Sciences 114:7061–7066.

Banglawala, N. 2010. Local adaptation under demographic and genetic fluctuations. (Doctoral dissertation). The University of Edinburgh.

Barton, N., and B. O. Bengtsson. 1986. The barrier to genetic exchange between hybri-dising populations. Heredity 57:357–376.

Barton, N., and A. Etheridge. 2018. Establishment in a new habitat by polygenic adaptation. Theoretical Population Biology 122:110–127.

Barton, N. H., and M. A. R. De Cara. 2009. The evolution of strong reproductive isolation. Evolution: International Journal of Organic Evolution 63:1171–1190.

Black, A. J., and A. J. McKane. 2012. Stochastic formulation of ecological models and their applications. Trends in Ecology & Evolution 27:337–345.

Blanquart, F., S. Gandon, and S. L. Nuismer. 2012. The effects of migration and drift on local adaptation to a heterogeneous environment. Journal of Evolutionary Biology 25:1351–1363.

Carroll, S. P., and C. Boyd. 1992. Host race radiation in the soapberry bug: Natural history with the history. Evolution 46:1052–1069.

Chevin, L.-M., O. Cotto, and J. Ashander. 2017. Stochastic evolutionary demography under a fluctuating optimum phenotype. The American Naturalist 190:786–802.

Coyne, J. A., N. H. Barton, and M. Turelli. 1997. Perspective: a critique of Sewall Wright’s shifting balance theory of evolution. Evolution 51:643–671.

Dobler, S., and B. Farrell. 1999. Host use evolution in Chrysochus milkweed beetles: evidence from behaviour, population genetics and phylogeny. Molecular Ecology 8:1297– 1307.

Fisher, R. A. 1922. On the mathematical foundations of theoretical statistics. Philosophical Transactions of the Royal Society of London. Series A, Containing Papers of a Mathematical or Physical Character 222:309–368.

———. 1930. The Genetical Theory of Natural Selection. Oxford: Clarendon Press.

Glover, K. A., M. F. Solberg, P. McGinnity, K. Hindar, E. Verspoor, M. W. Coulson, M. M. Hansen, H. Araki, Ø. Skaala, and T. Svåsand. 2017. Half a century of genetic interaction between farmed and wild Atlantic salmon: Status of knowledge and unanswered questions. Fish and Fisheries 18:890–927.

Gomulkiewicz, R., and R. D. Holt. 1995. When does evolution by natural selection prevent extinction? Evolution 49:201–207.

Gonzalez, A., O. Ronce, R. Ferriere, and M. E. Hochberg. 2013. Evolutionary rescue: an emerging focus at the intersection between ecology and evolution. Philosophical Transactions of the Royal Society B: Biological Sciences 368:20120404.

Govaert, L., E. A. Fronhofer, S. Lion, C. Eizaguirre, D. Bonte, M. Egas, A. P. Hendry, A. De Brito Martins, C. J. Melián, J. A. Raeymaekers, et al. 2019. Eco-evolutionary feedbacks – Theoretical models and perspectives. Functional Ecology 33:13–30.

Grant, P. R., and B. R. Grant. 2006. Evolution of character displacement in Darwin’s finches. Science 313:224–226.

Hanski, I., and T. Mononen. 2011. Eco-evolutionary dynamics of dispersal in spatially heterogeneous environments. Ecology Letters 14:1025–1034.

Kawecki, T. J. 2008. Adaptation to marginal habitats. Annual Review of Ecology, Evolution, and Systematics 39:321–342.

Kimura, M. 1955. Solution of a process of random genetic drift with a continuous model. Proceedings of the National Academy of Sciences of the United States of America 41:144.

Kinnison, M. T., and A. P. Hendry. 2001. The pace of modern life ii: From rates of contemporary microevolution to pattern and process. Genetica 112:145–164.

Kokko, H., and A. López-Sepulcre. 2007. The ecogenetic link between demography and evolution: can we bridge the gap between theory and data? Ecology Letters 10:773– 782.

Lande, R. 1993. Risks of population extinction from demographic and environmental stochasticity and random catastrophes. The American Naturalist 142:911–927.

Lande, R., S. Engen, and B.-E. Saether. 2003. Stochastic population dynamics in ecology and conservation. Oxford University Press, U.S.A.

Lande, R., S. Engen, and B.-E. Sther. 1998. Extinction times in finite metapopulation models with stochastic local dynamics. Oikos 83:383–389.

Lion, S. 2018. Theoretical approaches in evolutionary ecology: environmental feedback as a unifying perspective. The American Naturalist 191:21–44.

Mangel, M., and C. Tier. 1993. Dynamics of metapopulations with demographic stochasticity and environmental catastrophes. Theoretical Population Biology 44:1–31.

Ovaskainen, O., and B. Meerson. 2010. Stochastic models of population extinction. Trends in Ecology & Evolution 25:643 – 652.

Polechová, J. 2018. Is the sky the limit? on the expansion threshold of a species’ range. PLoS biology 16:e2005372.

Polechová, J., and N. H. Barton. 2015. Limits to adaptation along environmental gradients. Proceedings of the National Academy of Sciences 112:6401–6406.

Ronce, O., and M. Kirkpatrick. 2001. When sources become sinks: migrational meltdown in heterogeneous habitats. Evolution 55:1520–1531.

Rouhani, S., and N. Barton. 1993. Group selection and the shifting balance. Genetics Research 61:127–135.

Sachdeva, H. 2019. Effect of partial selfing and polygenic selection on establishment in a new habitat. Evolution 73:1729–1745.

Thompson, J. N. 1998. Rapid evolution as an ecological process. Trends in Ecology & Evolution 13:329–332.

Tufto, J. 2001. Effects of releasing maladapted individuals: a demographic-evolutionary model. The American Naturalist 158:331–340.

Whitlock, M. C., and N. H. Barton. 1997. The effective size of a subdivided population. Genetics 146:427–441.

Wright, S. 1932. The roles of mutation, inbreeding, crossbreeding, and selection in evolution. Proceedings of the Sixth International Congress of Genetics 1:356–366.

